# *Hex*-MASP for Mapping the Whole-tissue Spatial Proteome and the Intra-brain Distribution of Monoclonal Antibodies

**DOI:** 10.64898/2026.05.01.722341

**Authors:** Shihan Huo, Min Ma, Shuo Qian, Ming Zhang, Jie Pu, Xiaoyu Zhu, Sailee Rasam, Tara Barone, Robert Plunkett, Chi Zhou, Jun Qu

## Abstract

Whole-tissue spatial proteomics level provides critical insights into region-specific biological regulations but remains challenging. Previously, we introduced the Micro-scaffold Assisted Spatial Proteomics (MASP) concept for whole-tissue mapping. However, this prototype required substantial development in spatial resolution, practicality, and throughput for practical application. Here we present a next-generation MASP technique (*hex-*MASP) featuring i) a new design of hexagonal-micro-wells fabricated with optimized Projection Micro-Stereolithography (PµSL) 3D-printing, achieving high spatial resolution, sampling robustness and mechanical strength for reproducibly compartmentalizing even tough tissues; ii) enhanced throughput/effectiveness in sample preparation and LC-MS analysis with high quantitative quality. Applied to mouse brain, *hex-*MASP for the first time achieved in-depth, whole-tissue mapping for >6,000 proteins in mouse brains, with high spatial accuracy and excellent data quality. The substantially improved resolution revealed critical regional details across the entire brain, that were not previously captured, enabling precise depiction of protein distribution heterogeneity. This technique enabled the discovery of many unreported regionally-enriched proteins across brain structures. We further applied *hex-*MASP to investigate the intra-brain distribution of intracerebroventricularly-dosed antibody therapeutics and related proteins, which to our knowledge, enabled whole tissue mapping of protein drugs for the first time and revealed novel mechanistic insights into antibody distribution and localized treatment effects. *Hex*-MASP represent a robust, scalable platform for whole-tissue spatial proteomics.

## Introduction

Spatial proteomics at the whole-tissue level offers a panoramic view of the distribution of key proteins, such as regulators, biomarkers, enzymes, and receptors, across entire tissue sections. Such approach may provide novel insights into spatially-organized biological processes, enabling the integration of functional and pharmaceutical information with spatial information, and facilitating the detailed characterization of biologically and pharmaceutically meaningful region-to-region variability on the whole-tissue level^1^. For instance, the heterogeneous distribution of protein drugs, their targets, and biomarkers across different tissue regions can significantly impact therapeutic efficacy and toxicity^2, 3, 4, 5, 6, 7^. Therefore, understanding such distribution patterns is crucial for guiding therapeutic strategies and pharmaceutical development.

Currently, most existing spatial proteomics techniques are limited to the mapping of microscopic-scale tissue regions. For example, Laser Capture Microdissection (LCM)-based techniques involve sequential dissections a small number of microscopic tissue sections, followed by proteomics analysis using LC-MS^8, 9^. The Deep Visualization Proteomics (DVP) method allows for the dissection and analysis of individual cells or even subcellular components from tissues^10, 11^. While these groundbreaking approaches provide an excellent means to compare proteomic profiles between specific, microscopic regions, they are not suitable for whole-tissue level spatial analysis. The primary limitation lies in the sampling process, which procures micro-specimens one at a time, rendering the analysis of entire tissues impractically time-consuming and expensive.

To address this need, we previously developed Micro-scaffold Assisted Spatial Proteomics (MASP), which enables in-depth and accurate whole-tissue proteomics mapping^1^. This innovative concept, as shown in its prototype, utilizes a 3D-printed micro-scaffold in a rapid single-step cutting process, generating all location-specific micro-specimens across the entire tissue while precisely preserving their spatial context in tissue. The specimens are then analyzed via IonStar-based proteomics^12, 13, 14^, and the quantitative data are processed using the MAsP informatics package to reconstruct whole-tissue protein distribution maps.

However, to fully support the practical application of the MASP method and realize its potential, further critical development of this initial prototype is necessary, particularly in achieving higher spatial resolution, improved operational robustness of spatially-resolved micro-sampling, and enhanced reproducibility and throughput. Recognizing these needs, in this study we have developed a next-generation MASP technique, characterized by a new design of micro-scaffolds with substantially increased spatial resolution and operational robustness, as well as an efficient pipeline improving both quantitative reproducibility and throughput.

A key goal in this transformative upgrade was to improve spatial resolution, which enables more detailed and precise representations of protein distribution, thereby more accurately characterizing spatially-organized biological processes within tissues. To improve the resolution, high-quality fabrication of smaller micro-wells with sufficient strength to withstand the pressure needed for robust tissue compartmentalization is crucial. Our previous endeavors with square-shaped micro-wells limited the practical resolution of the MASP prototype at 400-µm^1^. Here, we introduced a new design of hexagon-shaped micro-wells, which, owing to their geometric advantages^15^ and the state-of-the-art Micro-Stereolithography (PµSL) technique specifically optimized for MASP, offers much greater mechanical strength and stability than the previous square counterparts, and achieves satisfactory performance under high pressure (**Figure 1a**). With an optimally-designed and precisely-controlled technical procedure, we demonstrated the robust production of micro-scaffolds with 100-180 µm resolution and successfully applied them in micro-compartmentalization of various types of tissues. Moreover, because hexagons require the least total wall length among common geometric tiling shapes^16^, this design maximizes the tissue surface area available for compartmentalization. To enhance throughput for analysis of the micro-specimens, we employed a new pipeline that combines automated sample preparation and multiplexed TMTpro protein quantification.

**Figure 1.**
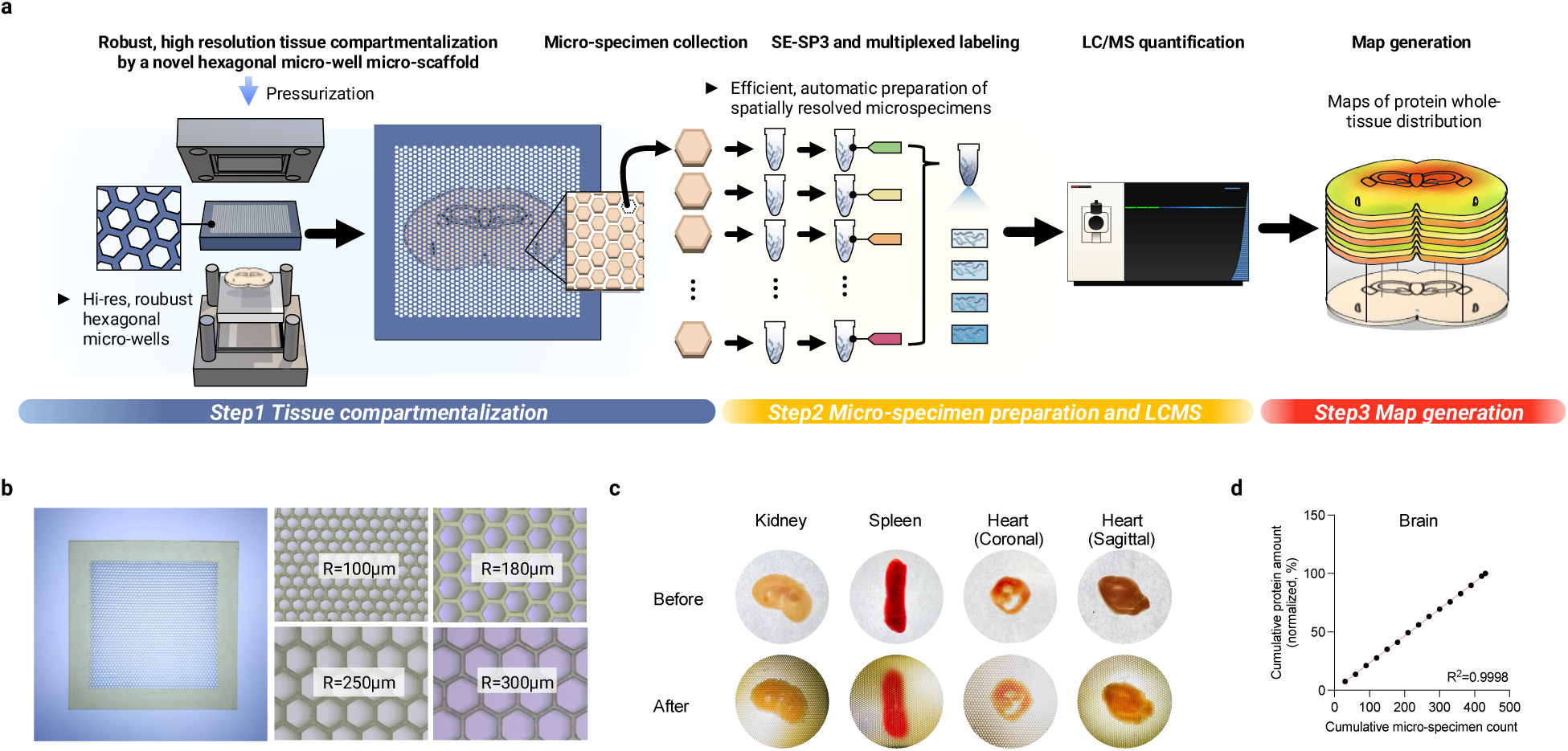
The next generation Micro-scaffold Assisted Spatial Proteomics (*hex*-MASP) pipeline with substantially higher spatial resolution, operational robustness and throughput. **(a)** Workflow overview (three major steps): Step 1 involves tissue compartmentalization via the newly engineered hexagonal micro-well micro-scaffolds, significantly improving spatial resolution and ensuring robust tissue compartmentalization and micro-specimen procurement. Step 2 is highly efficient, reproducible sample preparation using a µSE-SP3 sample preparation method developed here, followed by TMTpro labeling and LC-MS analysis. Step 3 generates and analyzes protein distribution maps using the MAsP module. **(b)** Examples of the new-generation hexagonal micro-scaffolds fabricated via optimized PµSL method, featuring substantially improved resolution, efficiency and mechanical strength. **(c)** Robust compartmentalization of most tissue types by the new-generation of micro-scaffolds, with precisely preserved spatial information and 100% sampling coverage. Mouse kidney, spleen and heart with coronal and sagittal views are shown as representative examples. **(d)** Excellent linearity (R² = 0.999) between cumulative protein amounts (%) and cumulative number of micro-specimens obtained from whole-brain compartmentalization, indicating robust and reproducible micro-sampling.

We demonstrated that this new generation of MASP pipeline (*hex*-MASP) enabled the whole-tissue-level spatial mapping of ∼6,000 proteins in mouse brain, providing substantially more detailed characteristics of the distribution of cerebral proteins, compared to the initial MASP prototype.

With this technique, we then explored the cerebral distribution of monoclonal antibody (mAb) drugs following intracerebroventricular (ICV) administration in mice. *Hex*-MASP allowed, for the first time, the mapping of the distribution of mAb across diverse brain regions, revealing the complex and heterogeneous nature of mAb penetration into the brain. Furthermore, facilitated by pattern analysis with the MAsP app, it was discovered that many cerebral proteins showed distributions that are positively- or negatively- correlated with these of the two mAbs. These discoveries provide novel insights into the mechanisms by which mAbs are distributed into brain post administration.

## Results

### Development of a next-generation micro-scaffold for practical and robust micro-compartmentalization of tissues with substantially enhanced spatial resolution

To enable mapping of proteins at the whole-tissue level, we previously developed a prototype termed Micro-scaffold Assisted Spatial Proteomics (MASP)^1^, consisting of three main steps: (1) tissue compartmentalization and procurement of location-specific micro-specimens; (2) micro-specimen sample preparation and subsequent LC-MS analysis; (3) generation of protein maps^1^. Step 1 is central to this strategy as it determines both the resolution and quality of protein mapping. Apparently, the ability to utilize smaller-size micro-wells for reliable tissue micro-compartmentalization is the key to improve spatial resolution but poses several technical challenges.

First, it is difficult for existing 3D printing protocols to fabricate precisely-spaced square micro-wells smaller than 400µm without compromising structural uniformity, which is vital for maintaining the spatial information during tissue compartmentalization process^1^. Second, the transition to smaller micro-wells requires significantly greater structural strength to withstand both the 3D fabrication process (including printing, rinsing, and curing) and the subsequent tissue compartmentalization step. Moreover, smaller wells increase mechanical resistance during tissue compartmentalization, necessitating higher pressure to ensure the complete separation of individual micro-specimens without structure collapse.

Previously, we demonstrated effective compartmentalization of brain tissue at a resolution of 400-µm with square-shaped micro-wells^1^. Nonetheless, attempts to enhance the micro-sampling resolution using the same fabrication techniques led to significantly reduced production yields, severe structural irregularities, as well as frequent failures in tissue compartmentalization, particularly in denser tissues like the heart and kidneys, owing to the structural collapse of the micro-wells (data not shown).

To overcome these limitations, here we developed a new-generation of micro-scaffolds, which achieves substantially improved spatial resolution for micro-compartmentalization of tissues and ensures high-quality protein mapping with precisely-reserved spatial information. We adopted two key approaches to realize these improvements: first, we replaced square-shaped micro-wells with the newly devised hexagonal micro-wells. Compared to the square micro-wells, the hexagon design inherently offers greater stability, leading to a high success rate of fabricating smaller micro-wells (detailed below) and enhanced mechanical strength, which is essential for sustaining the elevated pressure during the tissue compartmentalization with higher resolution. Moreover, hexagonal micro-wells provide superior space efficiency, reducing the total length of walls and thereby increasing the open space available for tissue compartmentalization. For example, if conditions are otherwise the same, hexagonal micro-wells provide a ∼14% more micro-well volume than using square micro-wells. Secondly, in MASP, it is essential to fabricate the micro-wells with near-perfect uniformity across the entire micro-scaffold, which poses a unique challenge for existing 3D printing techniques. We evaluated several state-of-the-art fabrication methods and selected the PµSL technique, which carries both high printing resolution and rapid layer-by-layer fabrication capabilities, rendering it a powerful technique to fabricate numerous precisely-spaced microstructures within a relatively large object^17^, as required here in printing the micro-scaffolds. Our pilot study found that comparing with other techniques, the PµSL produce micro-wells with smoother surface, more precise dimensions, and higher resolution.

To fabricate the hexagonal micro-well micro-scaffolds with high resolution, precision, and uniformity, we meticulously optimized the key parameters of the PµSL process. Here we highlight several key factors affecting the success of fabrication. First, we found that optimal energy exposure is critical, with different settings required for the first layer versus subsequent layers. After extensive evaluation, it was established that the UV light energy at 45 units and exposure time of 2s for the first layer, 1.6s for layers 2-5 and 0.8s for all subsequent layers achieved ideal results. Second, due to the unique design of numerous precisely-spaced, hollow micro-wells, parameters for the curing process are also crucial as it profoundly affects the mechanical strength and stability and uniformity of the scaffold device. After optimization, it was found that curing using ultraviolet (UV) light at 405 nm and at 60°C for 60 minutes, as well as a transient immobilization stage for 6 h produced high-quality scaffolds with excellent strength and minimal distortion. Finally, based on experimental optimization, we refined the design to achieve the best outcomes. For example, the ratio between the width of the outer frame and the size of the micro-wells was found to be important, as wider frames cause cracks in the micro-wells likely due to excessive bending stress, while frames too narrow lead to distorted micro-wells owing to the lack of structural stability needed to counteract the internal tension created during the printing process. It was found an optimal ratio of the side length of the micro-well zones vs. frame width provided a balanced mechanical stress distribution and enabled high success rate for production of high-quality scaffold. Optimal scaffold height (z-dimension) was also found crucial for achieving robust compartmentalization.

Utilizing these optimized conditions, we were able to fabricate high-resolution hexagonal micro-well micro-scaffolds with excellent uniformity and mechanical strength. Examples of scaffolds at 100, 180, 250 and 300 µm hexagons are shown in **Figure 1b**. The success rates for yielding perfectly uniform and crack-free micro-scaffolds were improved to >90% and >50% for 180-µm and 100-µm micro-scaffolds, respectively. These new generation of micro-scaffolds, with high mechanical strength and high spatial resolution, can efficiently compartmentalize most tissue types. **Figure 1c** shows the complete compartmentalization of brain, kidney, spleen and heart, with 100% coverage (*i.e.,* all tissue masses were turned to spatially-resolved micro-specimens without leftover) and without distortion of the original spatial information. Notably, even structurally challenging tissues like the heart, characterized by dense and muscular composition, were accurately and completely compartmentalized, which was not achievable with the previous MASP scaffold, even at a much lower resolution, due to the frequent distortion and collapse of micro-wells during this process.

The hexagonal scaffolds consistently yielded high-quality micro-specimens, demonstrated by exceptional linearity in cumulative protein amounts versus cumulative numbers of micro-specimens for brain tissue compartmentalized at a 180-µm resolution (R²=0.99, **Figure 1d**). This confirms the robust and reproducible micro-sampling capabilities provided by the next-generation hexagonal micro-well scaffolds.

### Optimization of a robust, high-throughput pipeline for sensitive proteomics quantification of the spatially-resolved micro-specimens

The improvement of spatial resolution necessitates the analysis of a larger number of micro-specimens, each with smaller sample sizes. This requires a high-throughput, automatic sample preparation method capable of sensitive proteomics quantification while maintaining high quantitative quality.

Previously, we demonstrated a micro-scale surfactant cocktail-aided extraction/precipitation/on-pellet digestion method (µ-SEPOD) for efficient preparation of micro-specimens in large cohorts^1^. One salient advantage of the µ-SEPOD method stems from its use of a high-concentration surfactant cocktail, achieving near-exhaustive tissue protein extraction, thorough protein denaturation for efficient digestion and effective removal of non-protein components^18^. However, its precipitation/cleanup step is difficult to automate, thereby limiting throughput.

To overcome this limitation, we aimed to develop an automated, high-throughput sample preparation strategy that retains the high recovery benefits provided by the surfactant cocktail. We combined the surfactant-cocktail extraction from µ-SEPOD with an automated clean-up and digestion procedure, known as the Single-pot, Solid-phase-enhanced Sample Preparation (SP3) approach^19, 20, 21^. The SP3 method, employing carboxylate-modified paramagnetic beads for protein enrichment, is well-suited for handling trace amounts of proteins in large cohorts of samples. Moreover, it is compatible with high surfactant concentrations, and adaptable to pipelines.

The key parameters of this new strategy, termed Micro Surfactant-cocktail Extraction with SP3 (µSE-SP3) have been thoroughly optimized to maximize recovery, reproducibility, and robustness. For example, we discovered that prolonged tissue extraction with sonication at 4°C with protease inhibitors markedly improved protein recovery and reproducibility (**Supplementary Figure 1a**) and observed both 50 mM Tris and 50 mM HEPES as digestion buffer were superior to 100 mM ammonium bicarbonate buffer which is recommended by prevailing SP3 protocols (**Figure 2a)**. HEPES digestion buffer was selected for its compatibility with subsequent TMTpro labeling.

**Figure 2.**
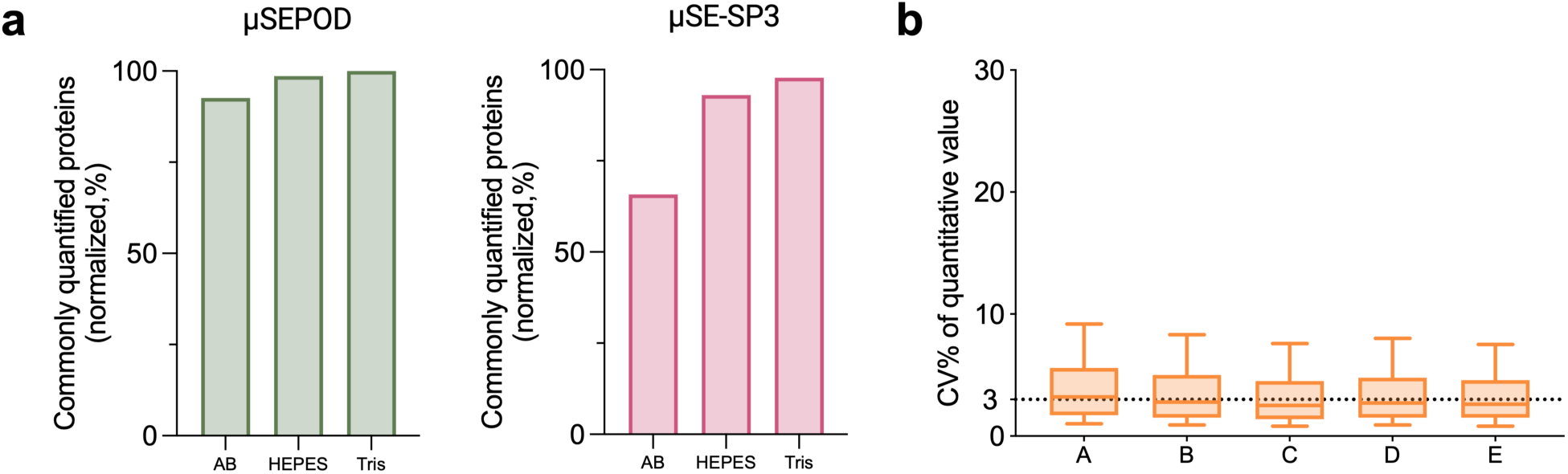
Development of sample preparation and LC-MS methods for micro-specimen proteomics quantification. **(a)** Comparison of proteome coverage achieved using µSEPOD and µSE-SP3 methods with different digestion buffers: Ammonium Bicarbonate (AB), Tris-HCl (Tris), and HEPES (n=3 per group). Values represent the number of proteins consistently quantified across all three replicates (normalized, %). **(b)** Under optimized conditions, multiplexed TMTpro labeling achieved low variability, indicated by a median coefficient of variation (CV%) of ∼3% (n=3 per group), demonstrating high analytical precision.

Comparative evaluations revealed that µSE-SP3 and µ-SEPOD yielded a similar number of quantifiable proteins from brain micro-specimens, while µSE-SP3 offered the significant advantage of automation. The median CV% of quantitative values among sample preparation replicates was only 8%, comparable to µ-SEPOD (7 %). The µSE-SP3 workflow exhibited high quantitative consistency in quantifying proteins (R^2^ =0.97-0.99, **Supplementary Figure 1b**) and high inter-plate and inter-day reproducibility (**Supplementary Figure 1c**), essential for analyzing the large-cohorts of micro-specimens.

To improve the throughput of LC-MS analysis, here we used Tandem Mass Tag Pro (TMTpro, 16-plex) approach to allow multiplexed quantification of up to 16 micro-specimens ^22^. Typically, TMT-based analysis requires extensive offline fractionation (12-24 fractions) for complex tissue samples to ensure the depth of analysis^23, 24, 25^, which diminishes its throughput benefits. Here, we explored the feasibility of using a smaller number of fractions for LC-MS analysis to strike a balance between achieving a reasonable analytical depth and improving throughput. Through rigorous optimization and evaluation on a sensitive and highly robust trapping-nano-LC-MS system^13^, we determined that using four concatenated fractions provides an optimal balance between throughput and depth of quantification. Details are in **Supplementary Results and Discussions**. We evaluated the quantitative performance of this optimized TMTpro-based approach against the IonStar method, previously utilized in MASP for accurate and in-depth proteomic quantification in large cohorts^1, 13, 14^, using a benchmark proteomic mixture of mouse, *E. coli*, and yeast (**Supplementary Figure 2a**; **Supplementary Results and Discussions**). At four concatenated fractions, this optimized TMTPro strategy achieved 80% of IonStar’s proteomics depth, while improving analytical throughput by four times (**Supplementary Figure 2b**). The median error% for quantification was <20% in all groups with excellent precision (median intra-group CV% were <3%, **Figure 2b**). Finally, we optimized the inter-batch normalization strategy (details in **Supplementary Results and Discussions**), effectively eliminating batch effects and achieved precise quantification across a large cohort of spatially specific micro-specimens (**Supplementary Figure 2c**).

### Assessing the accuracy of high-resolution protein mapping with the *hex*-MASP pipeline

We evaluated the mapping accuracy of *hex*-MASP using healthy mouse brain as the model system. Employing the *hex*-MASP pipeline, we successfully constructed distribution maps for ∼6,000 proteins across the mouse brain. The list of proteins that were mapped by *hex*-MASP is in **Supplementary Table 1**. The top10 GO Biological Processes of these mapped proteins are summarized in **Supplementary Table 2.** As illustrative examples, proteins maps involved in the Biological Processes with the greatest number of mapped proteins, protein transport, are shown in **Supplementary Figure 3a**. Biological Processes involved in phosphorylation, cell cycle, apoptotic process are shown in **Supplementary Figures 3b, 3c, and 3d**, respectively. As another example, the mapped proteins in KEGG pathways of neurodegeneration across multiple diseases are shown in **Supplementary Figure 3e**.

To evaluate the mapping accuracy of *hex*-MASP, we first surveyed the spatial distribution maps of known brain-region specific markers, such as the markers delineating the cortex, hypothalamus, thalamus, and striatum regions (**Figure 3a and Supplementary Figure 4a**). *hex*-MASP faithfully reproduced the expected regional distribution of these markers. Compared to the maps generated by the prototype version of MASP with squared micro-wells, the *hex*-MASP-derived maps displayed congruent overall patterns but with significantly enhanced details and clarity in demarcating brain regions.

**Figure 3.**
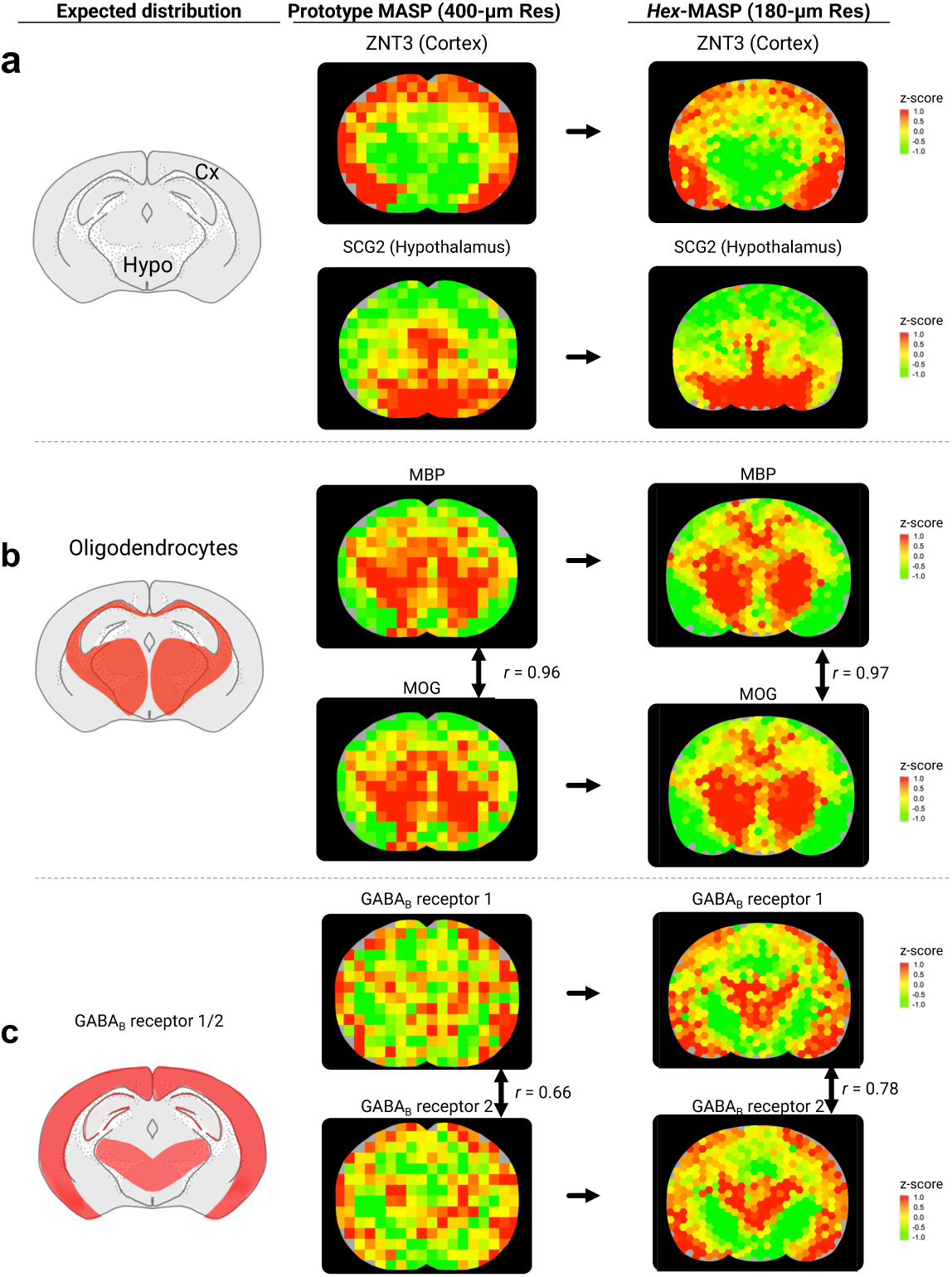
Mapping accuracy and resolution gain of *hex*-MASP (180 µm) versus the previous-generation MASP (400 µm). **(a)** Representative spatial maps for region-specific markers in cortex (Cx) and hypothalamus (Hypo). **(b)** Maps of well-known oligodendrocyte markers MBP and MOG show strong spatial correlation; **(c)** High spatial correlation between the two different proteins that form the same heterodimer, exemplified by GABA_B_ receptor subunits R1 and R2. Across panels, marker distributions match expected neuroanatomy and co-localized proteins exhibit high spatial correlation, supporting the mapping accuracy of *hex*-MASP. More importantly, the improved resolution of *hex*-MASP reveals critical structural details that were obscured in maps generated by the previous MASP technique.

In addition to the regional markers, we further examined whether *hex*-MASP could correctly depict the distribution of cell-type-specific markers within the brain, with the oligodendrocyte markers as examples. It was observed that *hex*-MASP accurately produced the expected distribution of well-known oligodendrocyte markers, such as MBP and MOG (**Figure 3b**). More examples of oligodendrocyte markers including CLD11, CN37 and MYPR, are presented in **Supplementary Figure 4b**. The maps of these oligodendrocyte markers showed remarkable similarity, with high correlation coefficients (Pearson r values of 0.96-0.99) when compared against MBP, confirming the high accuracy of *hex*-MASP’s spatial mapping. Moreover, comparing to the MASP prototype, *hex*-MASP provided significantly enhanced details, clearly revealing two well-defined, symmetric oligodendrocyte-dense regions in the inferior cerebral hemispheres, a delineation that was obscured by the previous version of MASP (**Figure 3b)**.

We further evaluated the mapping accuracy by comparing the distribution patterns of the two component proteins forming heterodimers. Given their totally different protein sequences, the two proteins in a heterodimer are independently mapped by *hex*-MASP; however, since these two proteins are co-localized in the dimmer form, their distribution maps should be highly correlated if the mapping method is accurate. Indeed, highly correlated distribution patterns of co-localized heterodimeric proteins were obtained by *hex*-MASP. For instance, the maps of GABA_B_ receptors 1 and 2 showed excellent correlation (**Figure 3c**). Again, it is clear that the higher resolution of *hex*-MASP provided more critical details than the MASP prototype. As shown in **Figure 3c**, for both the GABA_B_ receptors, the *hex*-MASP was able to define a heart-shaped central region of elevated receptor expression, complemented by peripheral ring-shaped distributions, which agrees well with the speculated distribution based on mRNA imaging data^26^. These intricate details were absent in the maps produced by the previous MASP^1^, demonstrating the superior spatial resolution offered by *hex*-MASP.

To summarize, the mapping accuracy of *hex*-MASP has been rigorously validated through the accurate reproduction of the anticipated distributions for known region-specific markers, the highly correlated distribution patterns among multiple markers for the same cell type, and the correct depiction of co-localization patterns of two heterodimeric protein components. Additionally, patterns observed in the *hex*-MASP generated maps were consistent with those in previously published maps by the prototype version of MASP, albeit with markedly more resolving power on spatial details, underscoring the methodological advancements and fidelity of *hex*-MASP in capturing the spatial proteomic landscape.

### Utility of *hex*-MASP for generating novel insights from a spatial dimension

*Hex*-MASP facilitated the accurate mapping of cerebral distributions for thousands of proteins at the whole-tissue level. This extensive dataset serves not only as a novel, comprehensive protein atlas of the brain but also significantly advances our understanding of the brain’s spatial architecture, offering new insights towards the spatially-organized biological processes. Here we highlight two specific application directions among numerous potential uses of *hex*-MASP-generated maps.

Firstly, *hex*-MASP enables the discovery of novel brain region specific and cell type specific protein markers. Utilizing the integrated MAsP app within the *hex*-MASP pipeline, we identified many cerebral proteins whose spatial distributions highly correlated with specific brain regions and the distribution of certain cell types. Importantly, many of these proteins’ regional expression patterns were previously unknown, underscoring *hex*-MASP’s utility in discovering novel potential markers. Examples of newly identified regionally-expressed proteins discovered in this study are illustrated in **Figure 4a**, including potential oligodendrocyte markers and proteins highly expressed in the cortex, hippocampus, and hypothalamus.

**Figure 4.**
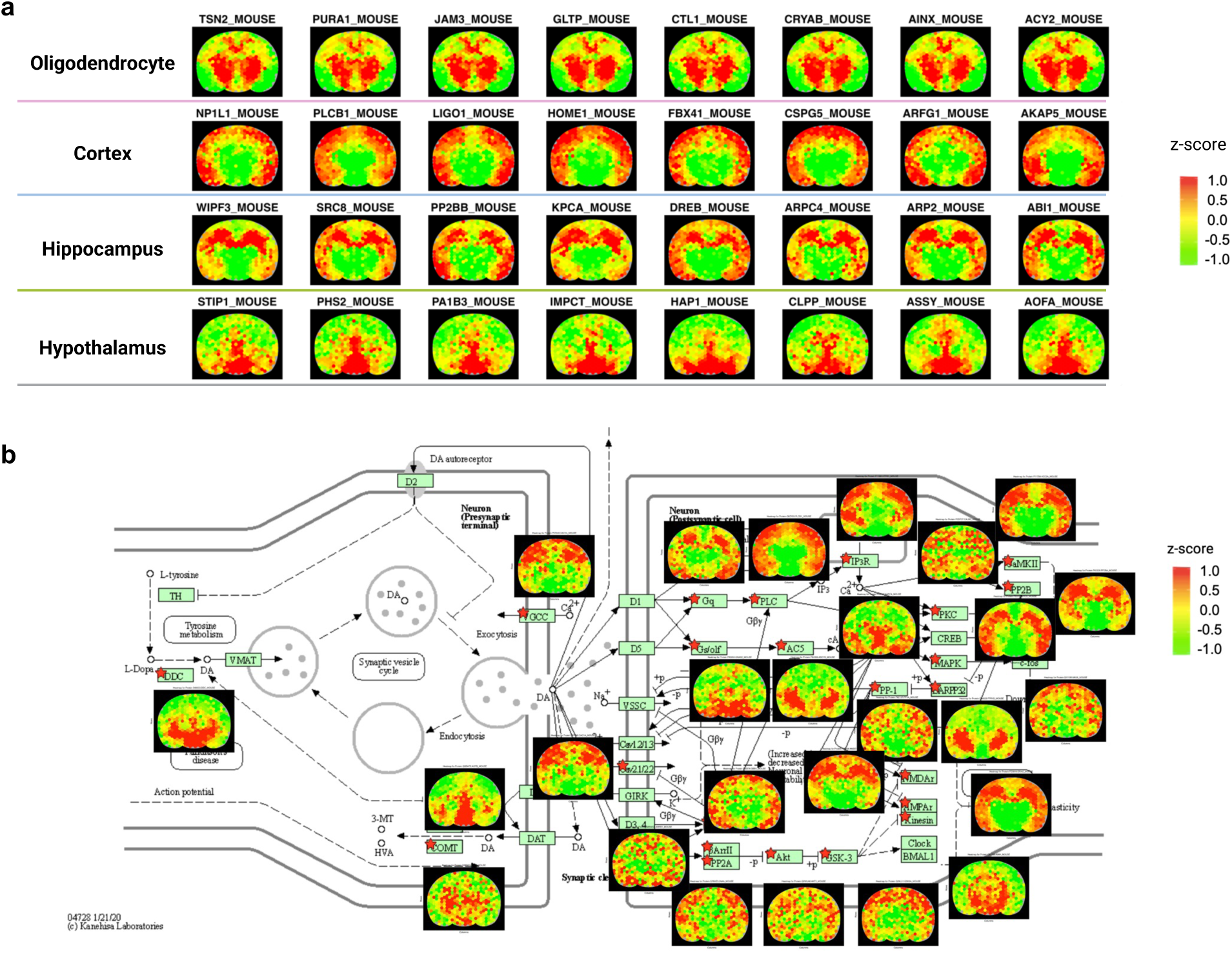
Application of *hex*-MASP for discovering novel region-specific and cell-type-specific markers, and for mapping the key proteins within biological pathways. **(a)** Representative distribution maps of newly discovered regionally-expressed proteins, including potential markers for oligodendrocytes and proteins highly expressed in the cortex, hippocampus, and hypothalamus. **(b)** Distribution maps of key proteins involved in the KEGG dopaminergic synapse pathway, with *hex*-MASP successfully generating whole-brain maps for 69% of these proteins.

Secondly, the comprehensive set of over 6,000 high-quality protein maps generated by *hex*-MASP opened an avenue for studying the whole-brain distributions of proteins within specific signaling pathways or biological processes. For example, the spatial distribution of proteins implicated in the dopaminergic synapse pathways, which controls a variety of key functions such as motivations, rewarding, learning and memory^27^, was depicted in **Figure 4b**. Using *hex*-MASP, 69% of proteins involved in dopaminergic synapse pathways were successfully mapped. The enhanced resolution provided by *hex*-MASP over its prototype version provided greater details in mapping key regulatory proteins, allowing more precise and informative correlations of distribution patterns among proteins involved in the same biological cascade. Another example is shown in **Supplementary Figure 5**, demonstrating the successful mapping of most proteins within the Alzheimer’s disease pathway. Moreover, compared to the previous version of MASP, the enhanced resolution of *hex*-MASP revealed greater details in the spatial organization of regulatory proteins.

### Application of *hex*-MASP for measuring monoclonal antibody (mAb) drug distribution in brain after intracerebroventricular (ICV) administration and unraveling proteins associated with mAb distribution and drug effects

To demonstrate the utility of the *hex*-MASP platform on whole-tissue mapping of protein therapeutics and associated regulatory proteins, we applied it to investigate mAb distribution in mouse brain post intracerebroventricular (ICV) injections (**Figure 5a**).

**Figure 5.**
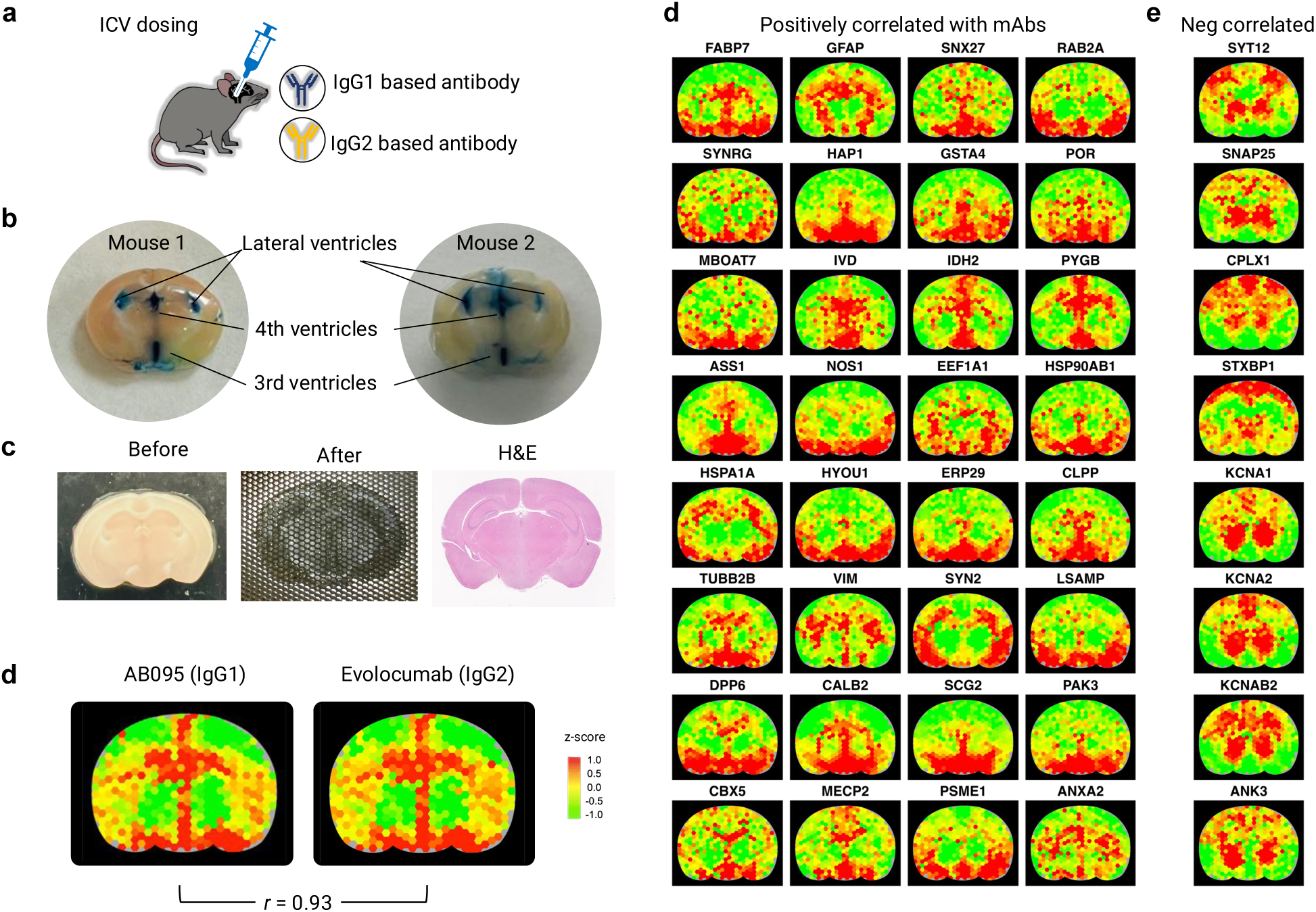
Application of *hex*-MASP to map whole-brain distribution of monoclonal antibodies (mAbs) and endogenous proteins following direct CNS administration. **(a)** Two mAbs, AB095 (IgG1) and Evolocumab (IgG2), were dosed to mice via Intracerebral ventricle (ICV) dosing. **(b)** Evans blue dye localized to the ventricular system without parenchymal spread across independent injections, indicating successful ICV delivery. **(c)** Photographs of mAbs-dosed brains before and after compartmentalization, together with H&E staining of an adjacent section, demonstrate precise preservation of spatial coordinates across the tissue. **(d)** Intra-brain distributions of the two ICV-dosed mAbs, quantified only using unique peptides, showed highly correlated patterns (r=0.93). **(e–f)** Examples of endogenous proteins whose Intra-brain distributions are positively correlated (**e,** r≥0.4, p<0.05) and negatively correlated (**f,** r ≤-0.4, p<0.05) to mAb distribution.

Antibody-based therapies show growing promise for the management of central nervous system disorders^28^. However, conventional administration routes such as intravenous, subcutaneous and intramuscular injections achieve limited brain exposure because the blood-brain barrier (BBB) restricts effective mAb delivery to target brain regions. ICV dosing has thus emerged as an important strategy to circumvent the BBB and thereby enhancing the delivery of antibody-based drugs into the brain. Previous studies suggested the heterogeneous distribution of antibodies within microscopically-discrete regions of the brain post an ICV administration^29^, however it remains unclear how ICV-dosed mAb differentially penetrate various regions of the brain. Studying the heterogeneous distribution of the mAb along with the thousands of cerebral proteins ***across the entire brain tissue***, will provide crucial information for understanding therapeutic efficacy and safety. For example, such study can identify which brain regions are more accessible to antibody drugs, elucidating mechanisms underlying heterogeneity of mAb distribution, as well as revealing spatially-organized biological functionality related to therapeutic effects and side effects in regions with antibody exposure. However, these investigations have traditionally been challenging largely due to the lack of a reliable, quantitative method for whole-tissue mapping.

Using *hex*-MASP, we measured the distributions of two types of mAb drugs along with thousands of cerebral proteins after a single ICV dosing, and its high-resolution capability enabled more accurate and precise distribution maps of mAbs compared to the prototype MASP. The purposes of study were: i) to characterize the heterogeneous mAb drug distribution across entire brain, thereby identifying brain regions with high mAb accessibility; ii) to discover the proteins whose spatial distribution patterns correlate with the administrated mAbs, either contributing to the mAb distribution (*e.g.,* by mediating mAb transport) or becoming dysregulated in regions exposed to mAbs.

The investigation was conducted in three steps: First, we developed and optimized an ICV administration protocol for murine brain, as detailed in **Experimental Section**. This protocol enabled reliable ICV injections despite the small size of mouse brain ventricles, as demonstrated by consistent Evans Blue dye distribution across the expected brain regions in independent injections, confirming successful ICV delivery (**Figure 5b)**.

Second, we performed a cassette ICV dosing of two mAbs, AB095 and Evolocumab, respectively of IgG1 and IgG2 types, each at 5mg/kg. The animals were euthanized 1-hour post-dosing, and brain tissues were harvested, sectioned, and compartmentalized by the *hex*-MASP procedure. Representative brain slice before and after compartmentalization, as well as H&E staining of an adjacent slice, are shown in **Figure 5c**. The compartmentalization process ***precisely preserved the spatial information in the tissue with full effectiveness*** (*i.e.,* all tissue mass was converted to spatially-resolved micro-specimens).

Third, with the developed pipeline described above, these micro-specimens were prepared, subjected to LC-MS analysis, and subsequently, distribution maps were generated using the MAsP app. The whole-tissue distribution maps of the two antibodies after ICV dosing are shown in **Figure 5d**. Interestingly, the maps of the two different types of IgGs exhibited highly correlated distribution patterns (r=0.93). Notably, both antibodies exhibited intensive distributions in periventricular regions, and showed decreasing levels outward into the parenchyma. These findings newly reveal the regional preference for ICV-administered antibodies.

To gain preliminary insights into the mechanisms underlying heterogeneous mAb penetration, we utilized the MAsP app to compute spatial correlations between the mAb distribution maps and >6,000 protein maps. Positive correlation was defined as r ≥0.4 and *p*<0.05, and negative correlation as r ≤-0.4 and *p*<0.05 to support functional exploratory analysis. In total, 99 proteins were positively correlated and 50 were negatively correlated with the mAb distribution (**Supplementary Table 3**). Representative examples are shown in **Figure 5e and 5f**. Many positively correlated proteins localize with regions of high CSF flow, intensive CSF-interstitial fluid exchange, and glymphatic activity, features facilitate mAb penetration after ICV dosing. For example, FABP7 and GFAP are markers of astrocytes, which scaffold CSF flow and perivascular transport^30, 31^.

Beyond anatomical co-localization, some positively correlated proteins point to homeostatic proteomic landscapes within brain regions exposed to antibodies. Here, we highlight several signaling pathways implicated by these proteins. It is well known that mAbs in tissues undergo intensive transcytosis via endosome trafficking, interacting with FcRn and Fcγ receptors (*e.g.,* on microglia)^32^. Consistent with this, proteins involved in vesicular trafficking and endosomal processes, such as SNX27, RAB2A, SYNRG, and HAP1^33, 34, 35^, showed maps aligned with mAb distribution.

Moreover, many of the proteins positively correlating with mAb distribution suggests an adaptive response to local IgG exposure: i) metabolic and redox enzymes involved in stress response (*e.g.,* GSTA4, POR, OAT, IVD, IDH2, PYGB, ASS1, NOS1), ii) protein synthesis/folding machinery, and chaperones (*e.g.,* EEF1A1, HSP90AB1, HSPA1A, HYOU1, ERP29, CLPP) for maintaining proteostasis; iii) cytoskeletal and structural proteins (e.g., TUBB2B, VIM, GFAP, SYN2, LSAMP) potentially contribute to neuron stabilization and iv) neuronal signaling and transport proteins (*e.g.,* DPP6, CALB2, SCG2, HAP1, PAK3). Finally, we observed that proteins associated with early immune response, such as PSME1 and ANXA2, showed spatially correlated expression with mAb distribution, which is consistent with previous findings that therapeutic antibodies can initiate early immune responses after tissue penetration^36^. Here, we observed that such a response could be elicited even by non-targeted IgG, suggesting that the Fc domain may activate immune surveillance mechanisms, which is supported by prior studies on Fc receptor-mediated responses^37^.

Conversely, many negatively correlated proteins were synaptic or neuron-specific markers (*e.g.,* SYT12, SNAP25, CPLX1, STXBP1, KCNA1, KCNA2). These markers are enriched in densely synaptic gray matter with high extracellular tortuosity and low extracellular volume fraction, conditions that impede diffusion of large molecules such as antibodies^38^. Similarly, markers associated with myelinated axons (*e.g.,* KCNAB2, ANK3) also showed negative correlations, consistent with reduced permeability in heavily myelinated regions^39^. Collectively, these above observations provide novel clues to direct future drug development efforts and generate interesting hypotheses warrants further investigation and validation.

## Discussion

Though the importance of spatial proteomics analysis has been widely acknowledged, currently available techniques for proteomic mapping at the whole-tissue level remain extremely limited. Our prior development of the MASP concept demonstrated the feasibility for comprehensive, in-depth mapping of thousands of proteins across entire tissue slices. Here, we introduced a substantially improved next-generation platform, *hex*-MASP, designed for practical applications by achieving crucial enhancements in spatial resolution, operational robustness, and analytical throughput. Central to *hex*-MASP is the implementation of newly designed micro-scaffolds featuring hexagonal micro-wells, optimized specifically for robust tissue compartmentalization with significantly improved spatial resolution and 100% sampling coverage. Compared to traditional square micro-wells, hexagonal designs with PµSL fabrication provides superior structural stability and spatial efficiency, achieving better mechanical strength and complete tissue transfer into micro-specimens. Additionally, it has been well-established in imaging science that hexagonal pixels offer advantages over square pixels, including improved accuracy, reduced directional bias, and enhanced inter-pixel connectivity that facilitates superior image processing outcomes such as noise filtering and pattern recognition^40^. These geometric advantages translate directly into enhanced proteomic mapping precision and quality in *hex*-MASP. Meanwhile, a robust sample preparation and LC-MS analysis strategy with significantly improved throughput was developed, including an automatic µSE-SP3 sample preparation leveraging a strong detergent cocktail for efficient and reproducible extraction, denaturation/cleanup and digestion, and multiplexed TMT-based quantification.

The high quantitative precision enables *hex*-MASP to sensitively detect differences of protein levels across various tissue regions. The mapping accuracy was rigorously validated via cross-referencing established region-specific brain markers and verifying consistent spatial patterns of protein heterodimers. *Hex*-MASP demonstrated substantial improvements over its predecessor by producing significantly higher-resolution maps capable of accurately characterizing finer details of protein distribution heterogeneity within tissues. These capabilities were exemplified through applications including the identification of novel brain-region-specific and cell-type-specific protein markers, and comprehensive mapping of spatial patterns of proteins involved in key signaling pathways. Moreover, we applied the pipeline in studying the intracerebral distribution of mAbs after a CNS dosing. Our results uncovered intricate spatial distribution patterns of the two most important types of therapeutic mAbs and identified multiple proteins exhibiting spatial distributions positively or negatively correlated with mAb penetration, implicating the roles of these proteins in the mediation of mAb distribution, therapeutic efficacy and safety. These results provided unprecedented insights into the spatial dynamics of mAb therapeutics penetration within the brain, and demonstrated the capacity of *hex*-MASP for studying the whole-tissue distribution of protein therapeutics and biomarkers, and for the comprehensive, spatially-resolved characterization of the signaling pathways and biological regulations associated with therapeutic efficacy and safety. Ongoing pharmaceutical validations aim to further substantiate these initial findings.

In summary, the new *hex*-MASP pipeline represents a significant advancement for whole-tissue, high-resolution spatial proteomics analysis. The micro-compartmentalization approach is compatible with various types of tissues, and the platform can be easily integrated with different sample preparation and analytical platforms. *hex*-MASP effectively addresses the prevailing challenges in accurate, in-depth whole-tissue proteomics mapping, thereby facilitating a better understanding of the complex spatial heterogeneity of proteins within tissues. The comprehensive datasets generated by *hex*-MASP, encompassing the maps of >6,000 proteins, provide a global perspective on biologically and pharmaceutically relevant regional variability and co-localized biological processes. These insights are valuable for generating testable hypotheses on spatially organized biological regulations, which can be further explored using the existing spatial proteomics techniques focusing on microscopic areas (*e.g.,* DVP or LMD) or antibody-based targeted strategies focusing on specific protein candidates.

## Methods

### Fabrication of the micro-scaffold and associated devices for reliable tissue compartmentalization

All 3D-fabricated devices used in the *hex*-MASP pipeline were designed using Autodesk Fusion 360 software (Autodesk Inc., USA). The new-generation micro-scaffolds featured precisely spaced hexagonal micro-wells with side lengths of 100, 180, 250, or 300 µm (**Figure 1b**). A custom 3D-printed pressurization module, comprising one upper and one lower components, enclosed a stacked assembly consisting of the micro-scaffold, tissue slice, and a polydimethylsiloxane (PDMS) supporting matrix, as illustrated in **Figure 1a**. Precisely regulated pressure, monitored by a Force-Sensing Resistor as described previously^1^, was applied to the top part of the module, which evenly transduced the pressure to the micro-scaffold and the supporting matrix to achieve complete, uniform micro-compartmentalization. The pressurization protocol involved two sequential stages: an initial lower pressure (∼4-7 kg) was applied for 10 seconds to immobilize the tissue, followed by a higher pressure (∼13-20 kg) to complete micro-compartmentalization. All micro-scaffolds and associated devices were printed using HTL-Y-20 resin on an ultra-high-resolution microArch S140 printer employing Projection Micro-Stereolithography (PµSL) technology (BMF, USA). Printing parameters, including layer exposure times, curing conditions, and detailed scaffold geometry, were rigorously optimized to ensure fabrication of the precisely-spaced high resolution hexagonal wells with exceptional uniformity and mechanical strength (see **Supplementary Experimental Procedure** for detailed conditions).

### The intracerebroventricular (ICV) dosing

All animal experiments performed in this study were conducted in accordance with the protocols approved by the Roswell Park Institutional Animal Care and Use Committee (IACUC). Healthy male C57BL/6J mice aged 10 weeks received intracerebroventricular (ICV) injections of a dosing cassette containing 5 mg/kg AB095 and 5 mg/kg Evolocumab. Injections were precisely administered using a stereotaxic frame (David Kopf Instruments, Tujunga, CA) at the following coordinates relative to bregma: +0.3 mm anterior, -0.8 mm lateral (right), and -2.5 mm ventral. At 1-hour post-injection, mice were perfused with heparinized saline. Subsequently, brains were collected and sectioned into 800 µm-thick coronal slices using steel blades with an Adult Mouse Brain Slicer Matrix (BSMAS001-1, Zivic Instruments, USA). Each tissue slice was mounted onto a PDMS supporting matrix and rapidly frozen at -80°C for 30 minutes prior to micro-compartmentalization.

### Compartmentalization and micro-specimen procurement

The frozen slice was precisely compartmentalized into spatially-resolved micro-specimens using the newly designed, 3D-printed hexagonal micro-scaffold, following these steps: first, the slice was mounted on a layer of PDMS supporting matrix which was securely settled into the holder on the bottom part of the pressurization module, followed by mounting a pre-cooled (in -20℃) micro-scaffold on the tissue slice. Second, the top part of the pressurization module (pre-cooled) is assembled to enclose the stack, and then pressure is applied to the top plate of the pressurization module via a steel press machine (VEVOR, USA) with a well-controlled pressure, and the pressure was monitored by a Fafeicy force-sensing resistor (Adafruit, USA) connected to a calibrated digital ohmmeter (Cen-Tech, USA). The entire operation was inside an enclosed chamber as described previously^1^. The micro-specimens were pushed out sequentially by a 280µm fused silica tubing and transferred to protein lo-binding tubes at -5 to 0 °C, ∼70-80% humidity, and saturated CO2 in the specimen-procurement chamber.

### Proteomic sample preparation

Micro-SE-SP3 was used for preparing high-quality samples for the spatial mapping. Protein extraction, reduction and alkylation was performed using the previously described micro-surfactant cocktail-aided extraction/precipitation/on-pellet digestion (µ-SEPOD)^1^. Briefly, exhaustive protein extraction was conducted by adding surfactant cocktail buffer (50 mM Tris-FA pH=8.5, 150 mM NaCl, 0.5% sodium deoxycholate, 2% IGEPAL CA-630, 2% SDS, cOmplete^TM^ Mini, EDTA-free Protease inhibitor tablets), followed by 15-min water bath sonication and 6-hr incubation at 4 °C. Five µg protein of each micro-specimen was reduced by 10 mM DTT (56 °C, 30min) and alkylated by 25 mM IAM in darkness (37°C, 30min). Protein cleanup and digestion were conducted using an automatic SP3 protocol on Kingfisher Flex (Thermo Fisher, Scientific, USA)^19, 20, 21^. The SP3 beads were added to the micro-specimen with a bead to protein ratio of 10:1. The protein was digested in 50mM HEPES (pH=8.5) at 37°C for 16 h with constant agitation. A reference sample containing a total of 200µg of proteins was pooled using aliquots from 40 randomly selected micro-specimen lysates and processed in parallel with the micro-specimen using the same protocol, for TMT inter-batch normalization. Details of TMTpro 16 plex labeling and off-line high-pH reversed-phase fractionation can be found in **Supplementary Experimental Procedure.**

### Liquid chromatography-mass spectrometry

A trapping-nano LC-high resolution MS system was used^12, 13^. The LC-MS system consists of an UltiMate 3000 gradient Micro LC system, an UltiMate 3000 Nano LC system, a WPS-3000 autosampler, and an Orbitrap Fusion Lumos Tribrid Mass Spectrometer (Thermo Fisher Scientific, USA). The peptides were firstly delivered onto a trapping column (C18, 5µm, 5 mm × 300 µm I.D.) at a flow rate of 10 µL/min with 1% B. Then the peptides were loaded onto a nano-LC column (C18, 2.5 µm, 65 cm × 75 µm I.D.) and eluted at a flow rate of 250 nL/min using a gradient of 4% to 11% B for 5 min; 11% to 32% B for 117 min; 32% to 50% B for 10 min; 50% to 97% B for 1 min, isocratic at 97% B for 17 min. Mobile phases A and B for the nano-LC were 0.1% FA in 2% ACN, and 0.1% FA in 88% ACN.

The data was collected (Xcalibur v 4.2.47, Thermo Fisher Scientific) in the positive mode. The MS1 spectra were acquired in the *m/z* range of 375-1500, with a resolution of 120,000 (FWHM@*m/z*=200). The maximum injection time for MS1 was 50 ms, automated gain control (AGC) target was 4E5. The dynamic exclusion was set to 60 s, and the mass tolerance was ± 10 ppm. MS2 scans were performed in the Ion Trap with CID fragmentation. The isolation window was set to 0.7 Da. The scan rate was set to Rapid. The collision energy was 32%. The maximum injection time for MS2 was 50 ms, with the AGC target at 1E4. MS3 scans were collected in the Orbitrap with HCD fragmentation using a resolution of 50,000. The Synchronous precursor selection was enabled, and the top 10 MS2 fragment ions will be included for MS3 fragmentation. The collision energy was 50%. The maximum injection time for MS3 was set to 105 ms with the AGC target at 1E5. Details of protein identification and quantification can be found in **Supplementary Experimental Procedure**.

### Generation of protein distribution maps by MAsP app

The previously developed MAsP app^1^, featuring a graphical user interface (GUI), was used to facilitate the rapid and efficient generation of protein distribution maps. The protein intensities file and location files were imported to the app for map generation and post-processing. A manual with detailed information on the MAsP app can be found at https://github.com/JunQu-Lab/MAsP. Z-score of protein abundances were calculated using the app embedded function and were used for all protein maps in figures. Additionally, the MAsP app allows for further analysis. For instance, the Pearson correlation coefficient, used for correlation calculations, is readily available within the app.

## Acknowledgements

This work was supported by NIH grants U01 DK137113, and a Center of Protein Therapeutics consortium grant. Data presented in this work were obtained via the University at Buffalo Center for Proteomics and Bioanalysis.

## Author contribution

S.H., M.M., S.Q., M.Z. and J.Q. conceptualized, designed and developed the *hex*-MASP pipeline. M.Z. and C.Z. designed and fabricated the 3D-printing products. S.H., M.M., M.Z., J.P., X.Z., S.R., T.R., R.P. developed and conducted the experiments. S.H., M.M. and S.Q. conducted data analysis. S.Q. developed and wrote the MAsP app. S.H., M.M. and J.Q. wrote and organized the manuscript.

## Author information

These authors contributed equally: Shihan Huo, Min Ma

## Corresponding author

Correspondence to Jun Qu.

## Data availability

All data presented in this manuscript are available as supplementary data files. The mass spectrometry proteomics data have been deposited to the ProteomeXchange Consortium via the PRIDE partner repository with the dataset identifier PXD068767.

## Code availability

The code included in MAsP app is available at https://github.com/JunQu-Lab/MAsP.

