## Supplementary_Information for "*Hex*-MASP for Mapping the Whole-tissue Spatial Proteome and the Intra-brain Distribution of Monoclonal Antibodies"

**Jun Qu, Ph.D.**

**Keywords:** MASP, in-depth spatial proteomics, whole tissue mapping, accurate protein distribution, mAb intra-brain distribution

#### Supplementary Figures

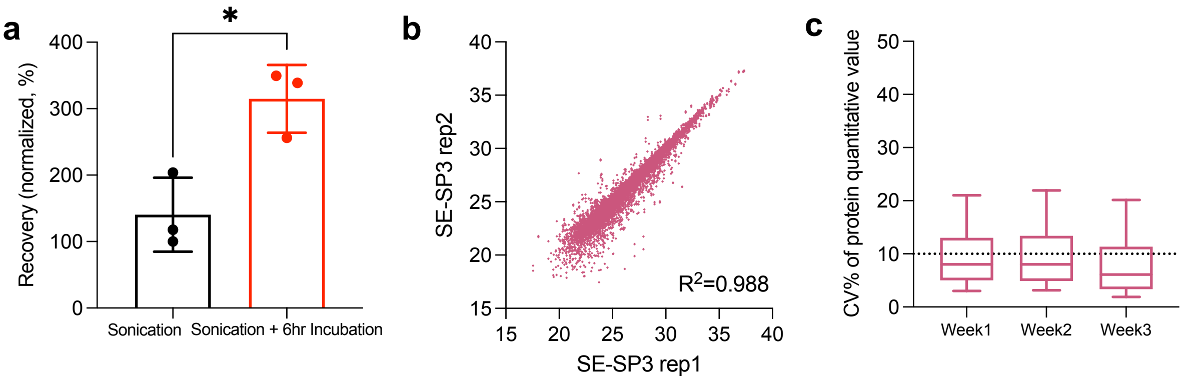

**Supplementary Figure 1. Optimization and evaluation of the μSE-SP3 sample preparation method.**

**(a)** Prolonged incubation (6 hours at 4°C) after sonication in the detergent cocktail significantly enhanced protein recovery from micro-specimens (collected from adjacent locations within the same brain region) compared to sonication alone (n=3 per group, $p$ = 0.016). **(b)** High correlation between log2 protein abundances obtained from two independent μSE-SP3 preparations using micro-specimens from adjacent locations within the same brain region, indicating strong quantitative consistency. **(c)** Excellent reproducibility of the μSE-SP3 method across plates and days, demonstrated by low coefficients of variation (CV%) in protein quantification over a three-week period using two separate plates.

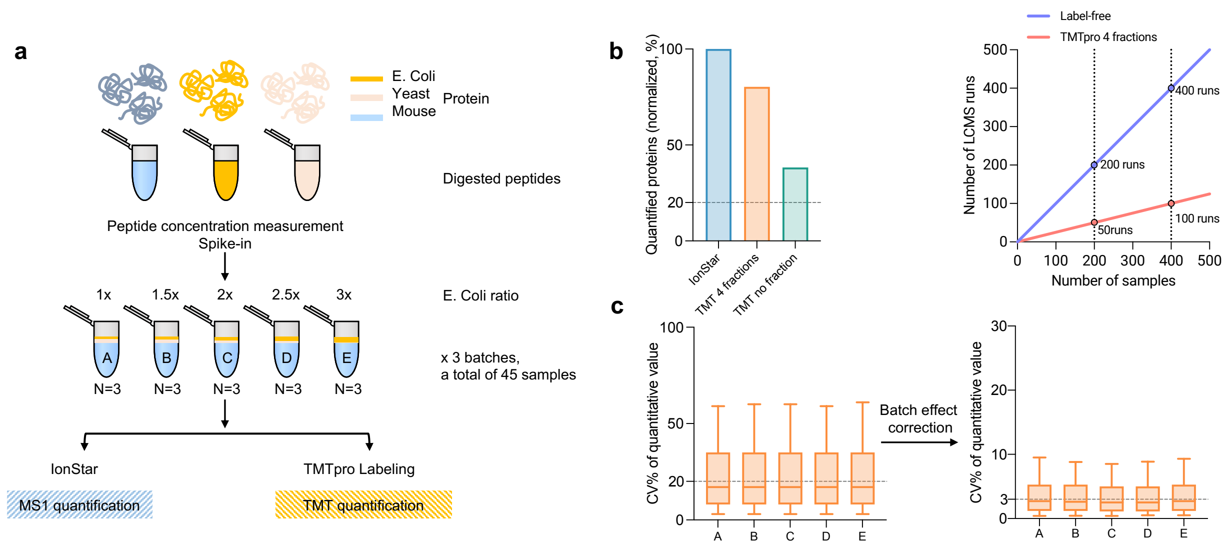

### Supplementary Figure 2. Experimental design and performance evaluation of the multiplexed, high-throughput LC-MS method.

**(a)** Scheme of experimental design. Benchmark mixed-proteome samples (Mouse, *E. coli*, and Yeast) containing variable amounts of *E. coli* protein were prepared in parallel as three sets (Groups A-E). One set was analyzed using IonStar (label-free, MS1-based quantification), while the other two sets were analyzed using TMTpro labeling approach. **(b)** Comparison of quantified proteins (normalized, %) between label-free Ionstar method and TMTpro with 4 fractions and no fractionation.(left) Schematic to compare of LC-MS throughput improvement achieved by using TMTpro labeling with four fractions relative to the IonStar label-free method.(right) **(c)** The quantitative method with normalization strategy maintained excellent quantitative reproducibility, as indicated by consistently low coefficients of variation (CV%) in protein quantification across two independent labeling batches (Groups A-E).

**a**

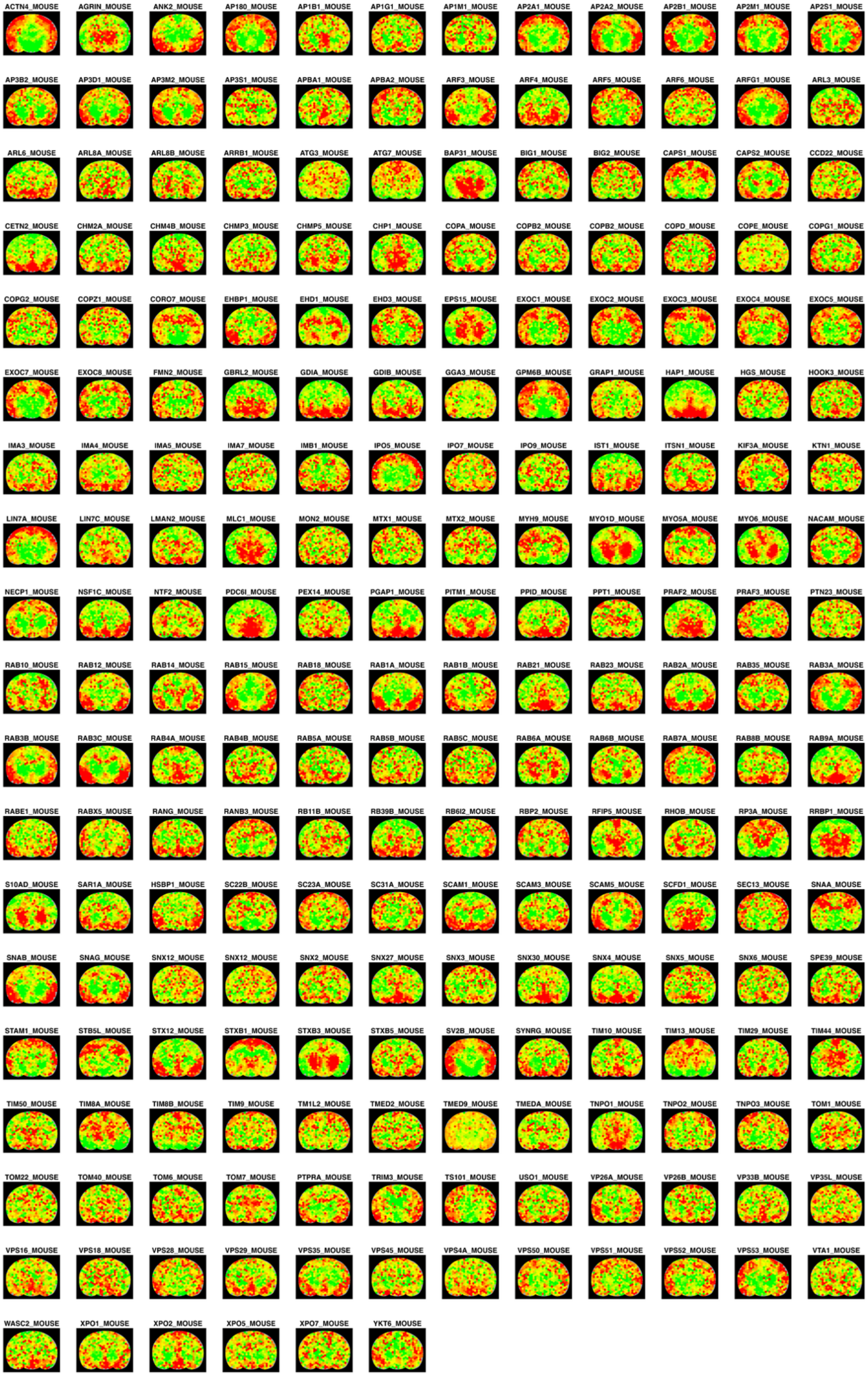
ACTN4_MOUSE
AGRIN_MOUSE
ANK2_MOUSE
AP180_MOUSE
AP1B1_MOUSE
AP1G1_MOUSE
AP1M1_MOUSE
AP2A1_MOUSE
AP2A2_MOUSE
AP2B1_MOUSE
AP2M1_MOUSE
AP2S1_MOUSE
AP3B2_MOUSE
AP3D1_MOUSE
AP3M2_MOUSE
AP3S1_MOUSE
APBA1_MOUSE
APBA2_MOUSE
ARF3_MOUSE
ARF4_MOUSE
ARF5_MOUSE
ARF6_MOUSE
ARFG1_MOUSE
ARL3_MOUSE
ARL6_MOUSE
ARL8A_MOUSE
ARL8B_MOUSE
ARRB1_MOUSE
ATG3_MOUSE
ATG7_MOUSE
BAP31_MOUSE
BIG1_MOUSE
BIG2_MOUSE
CAPS1_MOUSE
CAPS2_MOUSE
CCD22_MOUSE
CETN2_MOUSE
CHM2A_MOUSE
CHM4B_MOUSE
CHMP3_MOUSE
CHMP5_MOUSE
CHP1_MOUSE
COPA_MOUSE
COPB2_MOUSE
COPB2_MOUSE
COPD_MOUSE
COPE_MOUSE
COPG1_MOUSE
COPG2_MOUSE
COPZ1_MOUSE
CORO7_MOUSE
EHBP1_MOUSE
EHD1_MOUSE
EHD3_MOUSE
EPS15_MOUSE
EXOC1_MOUSE
EXOC2_MOUSE
EXOC3_MOUSE
EXOC4_MOUSE
EXOC5_MOUSE
EXOC7_MOUSE
EXOC8_MOUSE
FMN2_MOUSE
GBRL2_MOUSE
GDIA_MOUSE
GDIB_MOUSE
GGA3_MOUSE
GPM6B_MOUSE
GRAP1_MOUSE
HAP1_MOUSE
HGS_MOUSE
HOOK3_MOUSE
IMA3_MOUSE
IMA4_MOUSE
IMA5_MOUSE
IMA7_MOUSE
IMB1_MOUSE
IPO5_MOUSE
IPO7_MOUSE
IPO9_MOUSE
IST1_MOUSE
ITSN1_MOUSE
KIF3A_MOUSE
KTN1_MOUSE
LIN7A_MOUSE
LIN7C_MOUSE
LMAN2_MOUSE
MLC1_MOUSE
MON2_MOUSE
MTX1_MOUSE
MTX2_MOUSE
MYH9_MOUSE
MYO1D_MOUSE
MYO5A_MOUSE
MYO6_MOUSE
NACAM_MOUSE
NECP1_MOUSE
NSF1C_MOUSE
NTF2_MOUSE
PDC6I_MOUSE
PEX14_MOUSE
PGAP1_MOUSE
PITM1_MOUSE
PPID_MOUSE
PPT1_MOUSE
PRAF2_MOUSE
PRAF3_MOUSE
PTN23_MOUSE
RAB10_MOUSE
RAB12_MOUSE
RAB14_MOUSE
RAB15_MOUSE
RAB18_MOUSE
RAB1A_MOUSE
RAB1B_MOUSE
RAB21_MOUSE
RAB23_MOUSE
RAB2A_MOUSE
RAB35_MOUSE
RAB3A_MOUSE
RAB3B_MOUSE
RAB3C_MOUSE
RAB4A_MOUSE
RAB4B_MOUSE
RAB5A_MOUSE
RAB5B_MOUSE
RAB5C_MOUSE
RAB6A_MOUSE
RAB6B_MOUSE
RAB7A_MOUSE
RAB8B_MOUSE
RAB9A_MOUSE
RABE1_MOUSE
RABX5_MOUSE
RANG_MOUSE
RANB3_MOUSE
RB11B_MOUSE
RB39B_MOUSE
RB6I2_MOUSE
RBP2_MOUSE
RFIP5_MOUSE
RHOB_MOUSE
RP3A_MOUSE
RRBP1_MOUSE
S10AD_MOUSE
SAR1A_MOUSE
HSBP1_MOUSE
SC22B_MOUSE
SC23A_MOUSE
SC31A_MOUSE
SCAM1_MOUSE
SCAM3_MOUSE
SCAM5_MOUSE
SCFD1_MOUSE
SEC13_MOUSE
SNAA_MOUSE
SNAB_MOUSE
SNAG_MOUSE
SNX12_MOUSE
SNX12_MOUSE
SNX2_MOUSE
SNX27_MOUSE
SNX3_MOUSE
SNX30_MOUSE
SNX4_MOUSE
SNX5_MOUSE
SNX6_MOUSE
SPE39_MOUSE
STAM1_MOUSE
STB5L_MOUSE
STX12_MOUSE
STXB1_MOUSE
STXB3_MOUSE
STXB5_MOUSE
SV2B_MOUSE
SYNRG_MOUSE
TIM10_MOUSE
TIM13_MOUSE
TIM29_MOUSE
TIM44_MOUSE
TIM50_MOUSE
TIM8A_MOUSE
TIM8B_MOUSE
TIM9_MOUSE
TM1L2_MOUSE
TMED2_MOUSE
TMED9_MOUSE
TMEDA_MOUSE
TNPO1_MOUSE
TNPO2_MOUSE
TNPO3_MOUSE
TOM1_MOUSE
TOM22_MOUSE
TOM40_MOUSE
TOM6_MOUSE
TOM7_MOUSE
PTPRA_MOUSE
TRIM3_MOUSE
TS101_MOUSE
USO1_MOUSE
VP26A_MOUSE
VP26B_MOUSE
VP33B_MOUSE
VP35L_MOUSE
VPS16_MOUSE
VPS18_MOUSE
VPS28_MOUSE
VPS29_MOUSE
VPS35_MOUSE
VPS45_MOUSE
VPS4A_MOUSE
VPS50_MOUSE
VPS51_MOUSE
VPS52_MOUSE
VPS53_MOUSE
VTA1_MOUSE
WASC2_MOUSE
XPO1_MOUSE
XPO2_MOUSE
XPO5_MOUSE
XPO7_MOUSE
YKT6_MOUSE

**b**

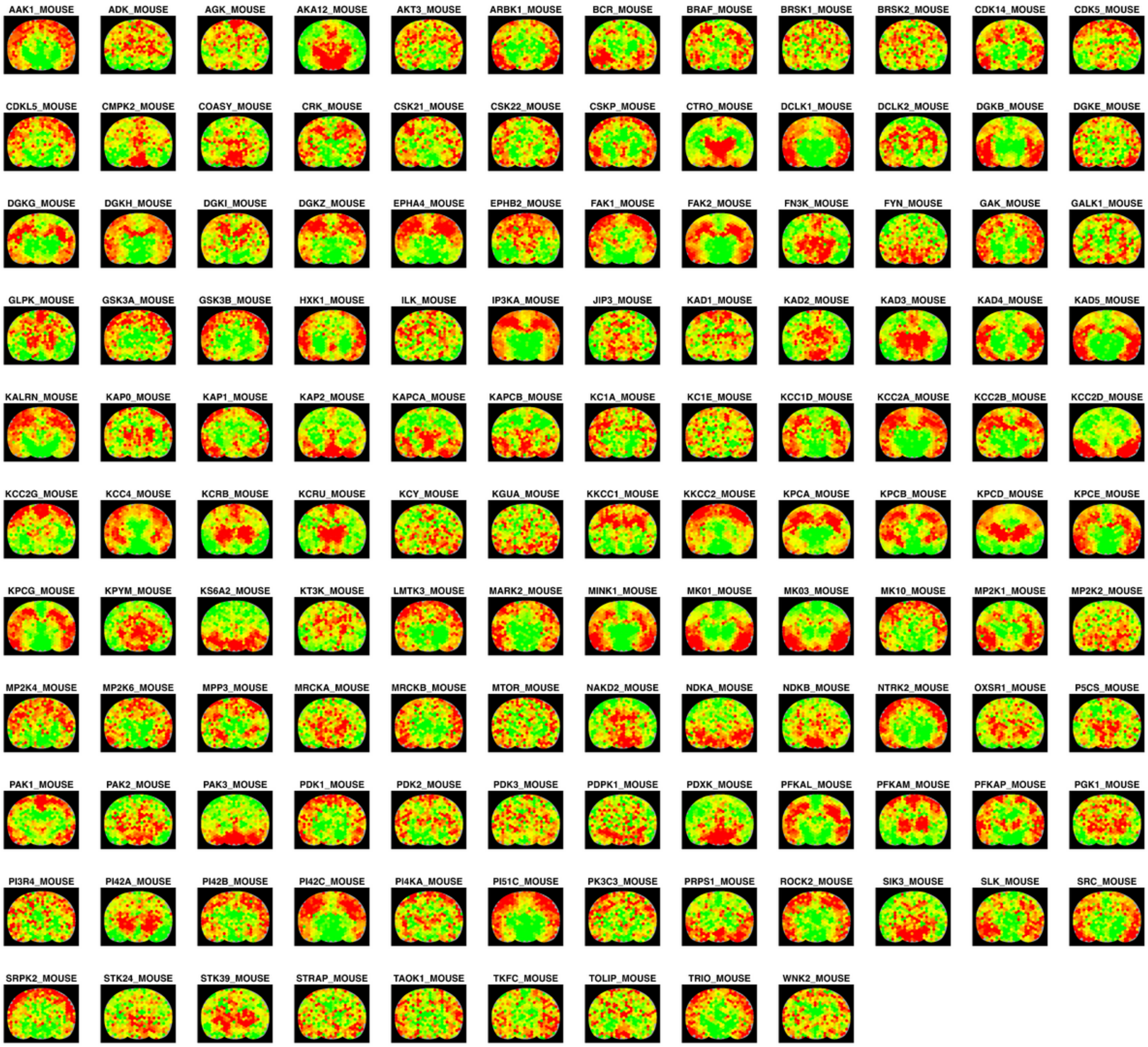
AAK1_MOUSE
ADK_MOUSE
AGK_MOUSE
AKA12_MOUSE
AKT3_MOUSE
ARBK1_MOUSE
BCR_MOUSE
BRAF_MOUSE
BRSK1_MOUSE
BRSK2_MOUSE
CDK14_MOUSE
CDK5_MOUSE
CDKL5_MOUSE
CMPK2_MOUSE
COASY_MOUSE
CRK_MOUSE
CSK21_MOUSE
CSK22_MOUSE
CSKP_MOUSE
CTRO_MOUSE
DCLK1_MOUSE
DCLK2_MOUSE
DGKB_MOUSE
DGKE_MOUSE
DGKG_MOUSE
DGKH_MOUSE
DGKI_MOUSE
DGKZ_MOUSE
EPHA4_MOUSE
EPHB2_MOUSE
FAK1_MOUSE
FAK2_MOUSE
FN3K_MOUSE
FYN_MOUSE
GAK_MOUSE
GALK1_MOUSE
GLPK_MOUSE
GSK3A_MOUSE
GSK3B_MOUSE
HXK1_MOUSE
ILK_MOUSE
IP3KA_MOUSE
JIP3_MOUSE
KAD1_MOUSE
KAD2_MOUSE
KAD3_MOUSE
KAD4_MOUSE
KAD5_MOUSE
KALRN_MOUSE
KAP0_MOUSE
KAP1_MOUSE
KAP2_MOUSE
KAPCA_MOUSE
KAPCB_MOUSE
KC1A_MOUSE
KC1E_MOUSE
KCC1D_MOUSE
KCC2A_MOUSE
KCC2B_MOUSE
KCC2D_MOUSE
KCC2G_MOUSE
KCC4_MOUSE
KCRB_MOUSE
KCRU_MOUSE
KCY_MOUSE
KGUA_MOUSE
KKCC1_MOUSE
KKCC2_MOUSE
KPCA_MOUSE
KPCB_MOUSE
KPCD_MOUSE
KPCE_MOUSE
KPCG_MOUSE
KPYM_MOUSE
KS6A2_MOUSE
KT3K_MOUSE
LMTK3_MOUSE
MARK2_MOUSE
MINK1_MOUSE
MK01_MOUSE
MK03_MOUSE
MK10_MOUSE
MP2K1_MOUSE
MP2K2_MOUSE
MP2K4_MOUSE
MP2K6_MOUSE
MPP3_MOUSE
MRCKA_MOUSE
MRCKB_MOUSE
MTOR_MOUSE
NAKD2_MOUSE
NDKA_MOUSE
NDKB_MOUSE
NTRK2_MOUSE
OXSR1_MOUSE
P5CS_MOUSE
PAK1_MOUSE
PAK2_MOUSE
PAK3_MOUSE
PDK1_MOUSE
PDK2_MOUSE
PDK3_MOUSE
PDPK1_MOUSE
PDXK_MOUSE
PFKAL_MOUSE
PFKAM_MOUSE
PFKAP_MOUSE
PGK1_MOUSE
PI3R4_MOUSE
PI42A_MOUSE
PI42B_MOUSE
PI42C_MOUSE
PI4KA_MOUSE
PI51C_MOUSE
PK3C3_MOUSE
PRPS1_MOUSE
ROCK2_MOUSE
SIK3_MOUSE
SLK_MOUSE
SRC_MOUSE
SRPK2_MOUSE
STK24_MOUSE
STK39_MOUSE
STRAP_MOUSE
TAOK1_MOUSE
TKFC_MOUSE
TOLIP_MOUSE
TRIO_MOUSE
WNK2_MOUSE

c

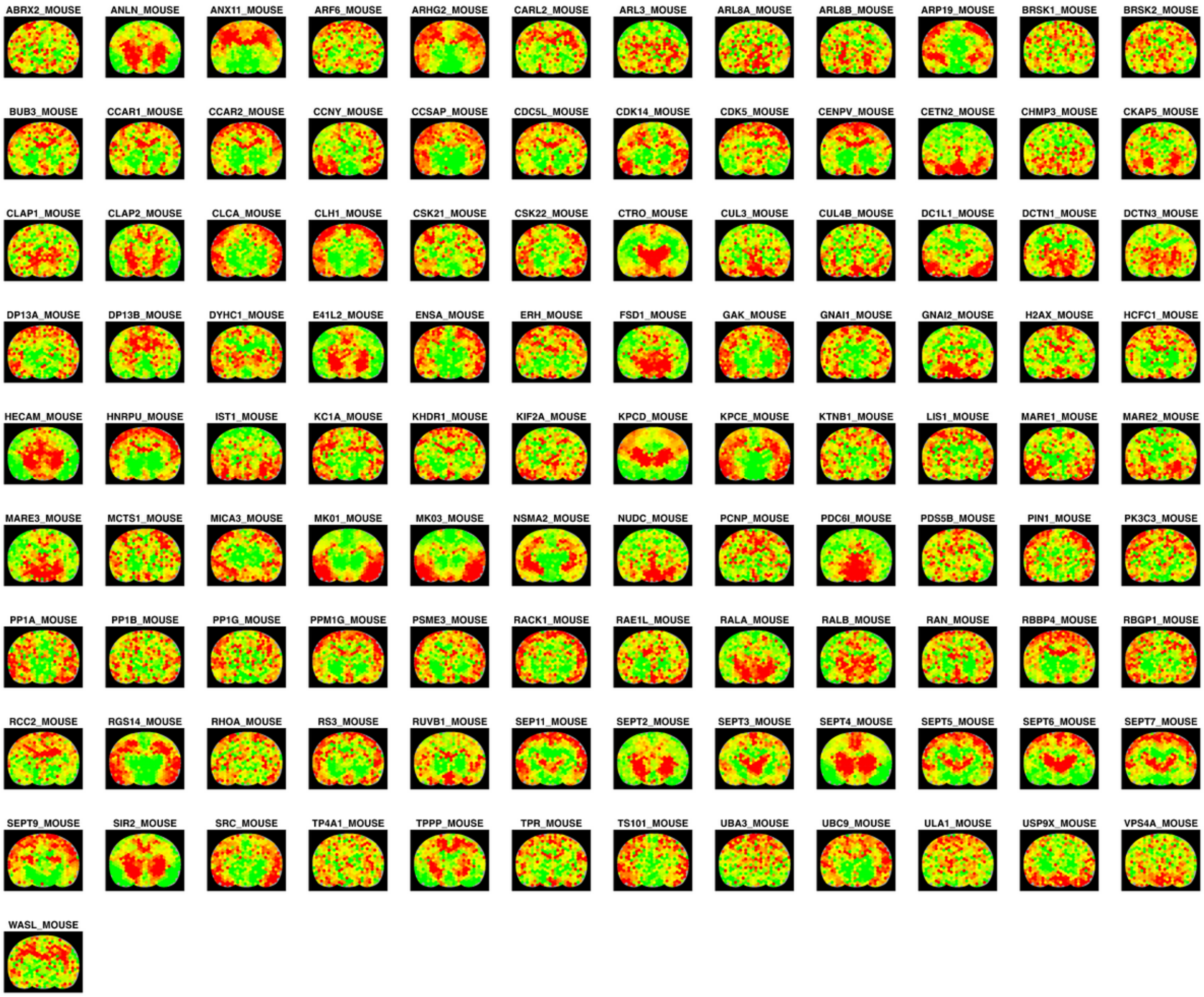
ABRX2_MOUSE
ANLN_MOUSE
ANX11_MOUSE
ARF6_MOUSE
ARHG2_MOUSE
CARL2_MOUSE
ARL3_MOUSE
ARL8A_MOUSE
ARL8B_MOUSE
ARP19_MOUSE
BRSK1_MOUSE
BRSK2_MOUSE
BUB3_MOUSE
CCAR1_MOUSE
CCAR2_MOUSE
CCNY_MOUSE
CCSAP_MOUSE
CDC5L_MOUSE
CDK14_MOUSE
CDK5_MOUSE
CENPV_MOUSE
CETN2_MOUSE
CHMP3_MOUSE
CKAP5_MOUSE
CLAP1_MOUSE
CLAP2_MOUSE
CLCA_MOUSE
CLH1_MOUSE
CSK21_MOUSE
CSK22_MOUSE
CTRO_MOUSE
CUL3_MOUSE
CUL4B_MOUSE
DC1L1_MOUSE
DCTN1_MOUSE
DCTN3_MOUSE
DP13A_MOUSE
DP13B_MOUSE
DYHC1_MOUSE
E41L2_MOUSE
ENSA_MOUSE
ERH_MOUSE
FSD1_MOUSE
GAK_MOUSE
GNAI1_MOUSE
GNAI2_MOUSE
H2AX_MOUSE
HCFC1_MOUSE
HECAM_MOUSE
HNRPU_MOUSE
IST1_MOUSE
KC1A_MOUSE
KHDR1_MOUSE
KIF2A_MOUSE
KPCD_MOUSE
KPCE_MOUSE
KTNB1_MOUSE
LIS1_MOUSE
MARE1_MOUSE
MARE2_MOUSE
MARE3_MOUSE
MCTS1_MOUSE
MICA3_MOUSE
MK01_MOUSE
MK03_MOUSE
NSMA2_MOUSE
NUDC_MOUSE
PCNP_MOUSE
PDC6I_MOUSE
PDS5B_MOUSE
PIN1_MOUSE
PK3C3_MOUSE
PP1A_MOUSE
PP1B_MOUSE
PP1G_MOUSE
PPM1G_MOUSE
PSME3_MOUSE
RACK1_MOUSE
RAE1L_MOUSE
RALA_MOUSE
RALB_MOUSE
RAN_MOUSE
RBBP4_MOUSE
RBGP1_MOUSE
RCC2_MOUSE
RGS14_MOUSE
RHOA_MOUSE
RS3_MOUSE
RUVB1_MOUSE
SEP11_MOUSE
SEPT2_MOUSE
SEPT3_MOUSE
SEPT4_MOUSE
SEPT5_MOUSE
SEPT6_MOUSE
SEPT7_MOUSE
SEPT9_MOUSE
SIR2_MOUSE
SRC_MOUSE
TP4A1_MOUSE
TPPP_MOUSE
TPR_MOUSE
TS101_MOUSE
UBA3_MOUSE
UBC9_MOUSE
ULA1_MOUSE
USP9X_MOUSE
VPS4A_MOUSE
WASL_MOUSE

**d**

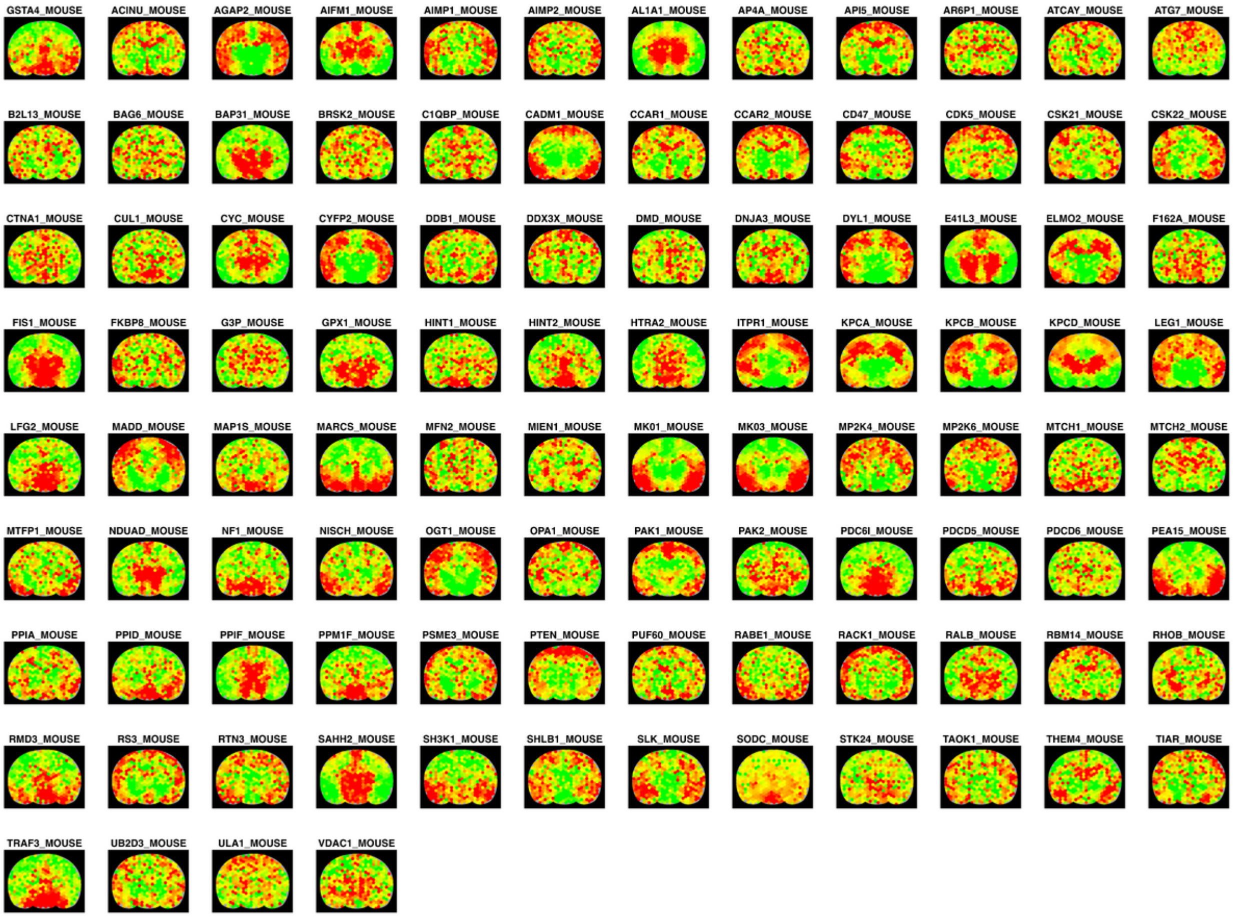
GSTA4_MOUSE
ACINU_MOUSE
AGAP2_MOUSE
AIFM1_MOUSE
AIMP1_MOUSE
AIMP2_MOUSE
AL1A1_MOUSE
AP4A_MOUSE
API5_MOUSE
AR6P1_MOUSE
ATCAY_MOUSE
ATG7_MOUSE
B2L13_MOUSE
BAG6_MOUSE
BAP31_MOUSE
BRSK2_MOUSE
C1QBP_MOUSE
CADM1_MOUSE
CCAR1_MOUSE
CCAR2_MOUSE
CD47_MOUSE
CDK5_MOUSE
CSK21_MOUSE
CSK22_MOUSE
CTNA1_MOUSE
CUL1_MOUSE
CYC_MOUSE
CYFP2_MOUSE
DDB1_MOUSE
DDX3X_MOUSE
DMD_MOUSE
DNJA3_MOUSE
DYL1_MOUSE
E41L3_MOUSE
ELMO2_MOUSE
F162A_MOUSE
FIS1_MOUSE
FKBP8_MOUSE
G3P_MOUSE
GPX1_MOUSE
HINT1_MOUSE
HINT2_MOUSE
HTRA2_MOUSE
ITPR1_MOUSE
KPCA_MOUSE
KPCB_MOUSE
KPCD_MOUSE
LEG1_MOUSE
LFG2_MOUSE
MADD_MOUSE
MAP1S_MOUSE
MARCS_MOUSE
MFN2_MOUSE
MIEN1_MOUSE
MK01_MOUSE
MK03_MOUSE
MP2K4_MOUSE
MP2K6_MOUSE
MTCH1_MOUSE
MTCH2_MOUSE
MTFP1_MOUSE
NDUAD_MOUSE
NF1_MOUSE
NISCH_MOUSE
OGT1_MOUSE
OPA1_MOUSE
PAK1_MOUSE
PAK2_MOUSE
PDC6I_MOUSE
PDCD5_MOUSE
PDCD6_MOUSE
PEA15_MOUSE
PPIA_MOUSE
PPID_MOUSE
PPIF_MOUSE
PPM1F_MOUSE
PSME3_MOUSE
PTEN_MOUSE
PUF60_MOUSE
RABE1_MOUSE
RACK1_MOUSE
RALB_MOUSE
RBM14_MOUSE
RHOB_MOUSE
RMD3_MOUSE
RS3_MOUSE
RTN3_MOUSE
SAHH2_MOUSE
SH3K1_MOUSE
SHLB1_MOUSE
SLK_MOUSE
SODC_MOUSE
STK24_MOUSE
TAOK1_MOUSE
THEM4_MOUSE
TIAR_MOUSE
TRAF3_MOUSE
UB2D3_MOUSE
ULA1_MOUSE
VDAC1_MOUSE

**e**

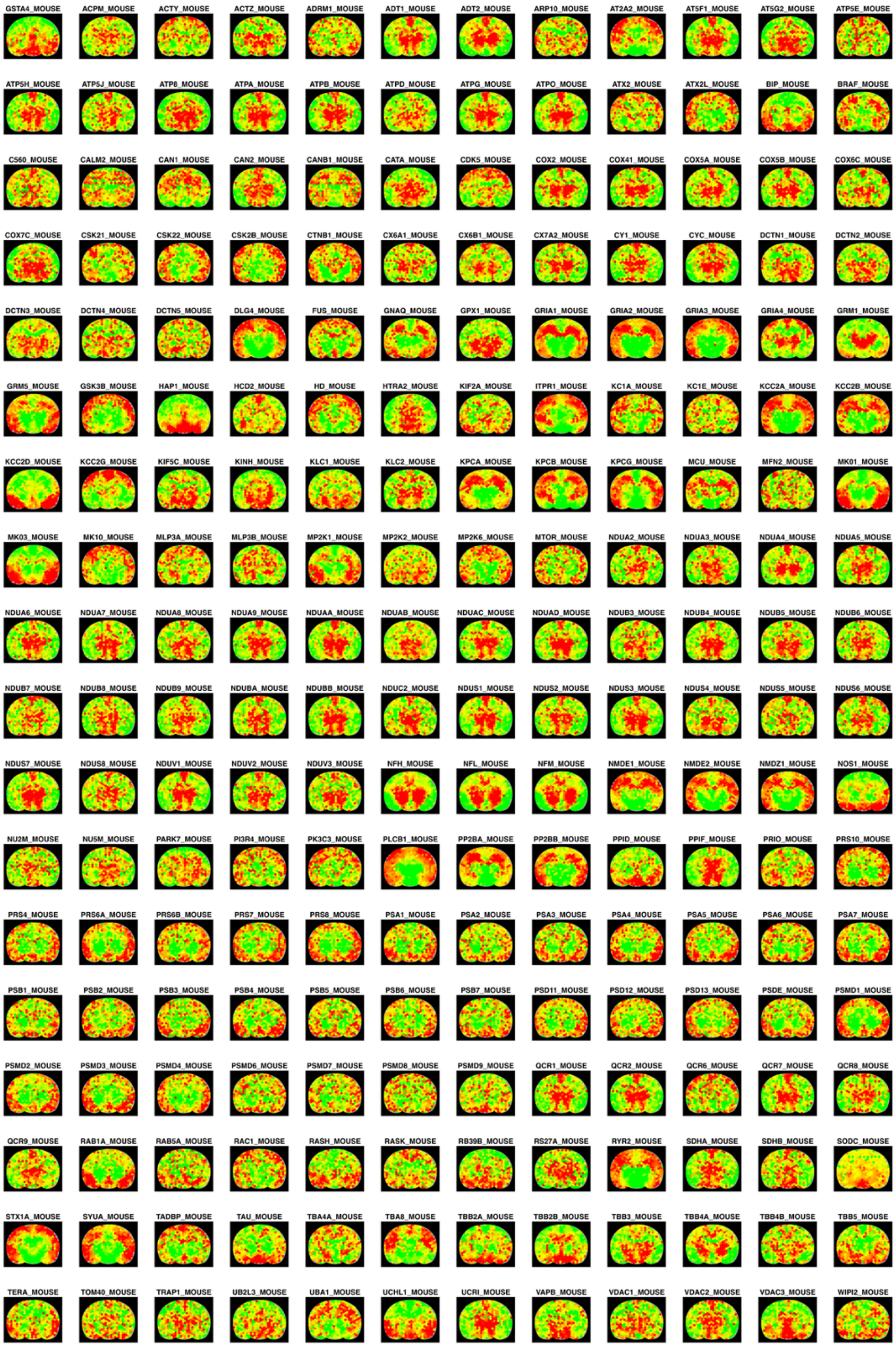

**Supplementary Figure 3. The maps of proteins involved in GO Biological Processes and KEGG pathways acquired by *Hex*-MASP.**

**(a)** GOBP protein transport, **(b)** GOBP phosphorylation, **(c)** GOBP cell cycle, **(d)** GOBP apoptotic process, **(e)** KEGG pathways of neurodegeneration-multiple diseases. The z-score color scale is from -1.0 (green) to 1.0 (red).

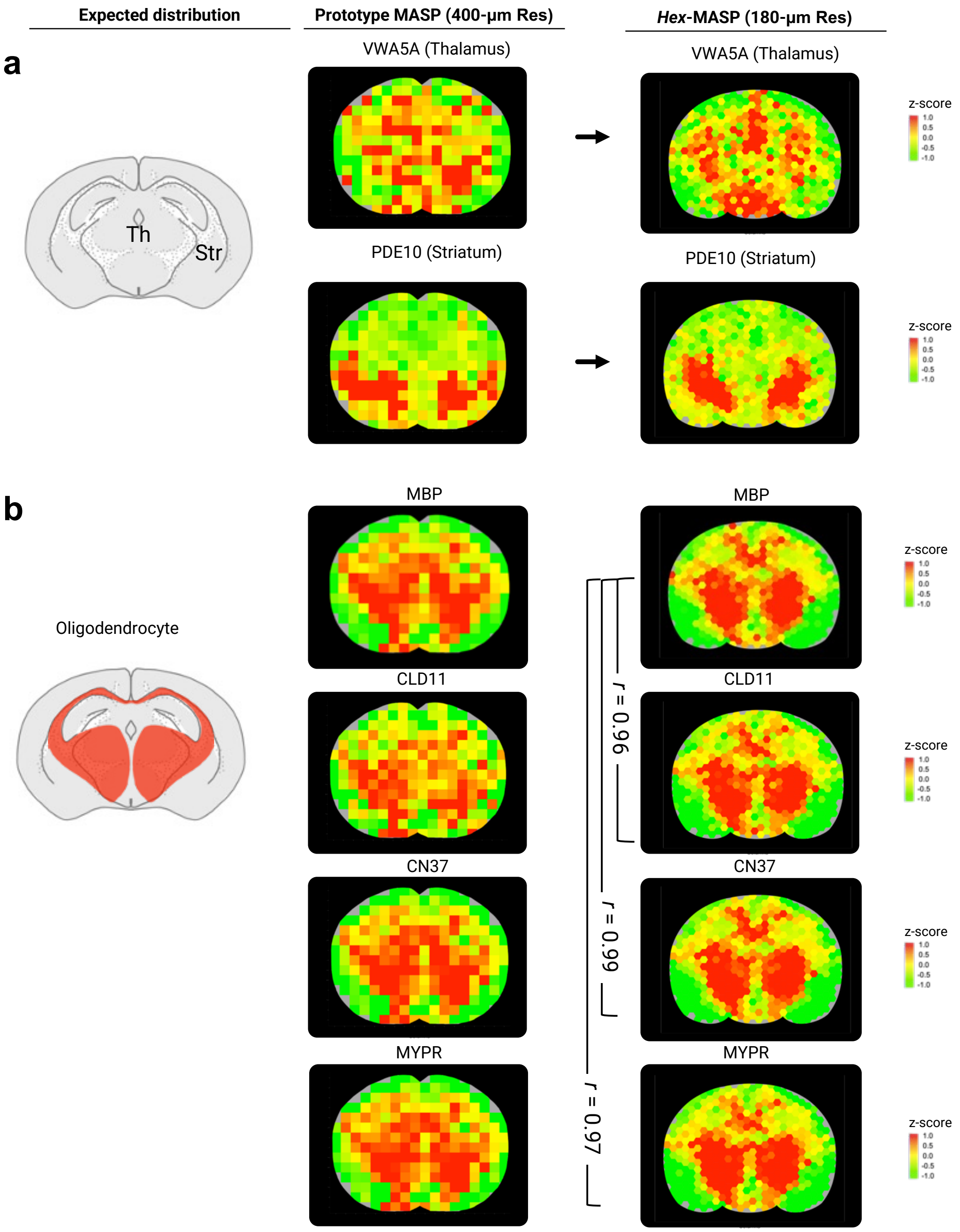

**Supplementary Figure 4. Validation of the high accuracy of protein maps generated using *Hex*-MASP (180 µm spatial resolution) and comparison with the previous generation MASP (400 µm spatial resolution):**

**(a)** Additional regional markers (Thalamus-Th and Striatum-St) and **(b)** Additional oligodendrocyte makers. The z-score color scale is from -1.0 (green) to 1.0 (red).

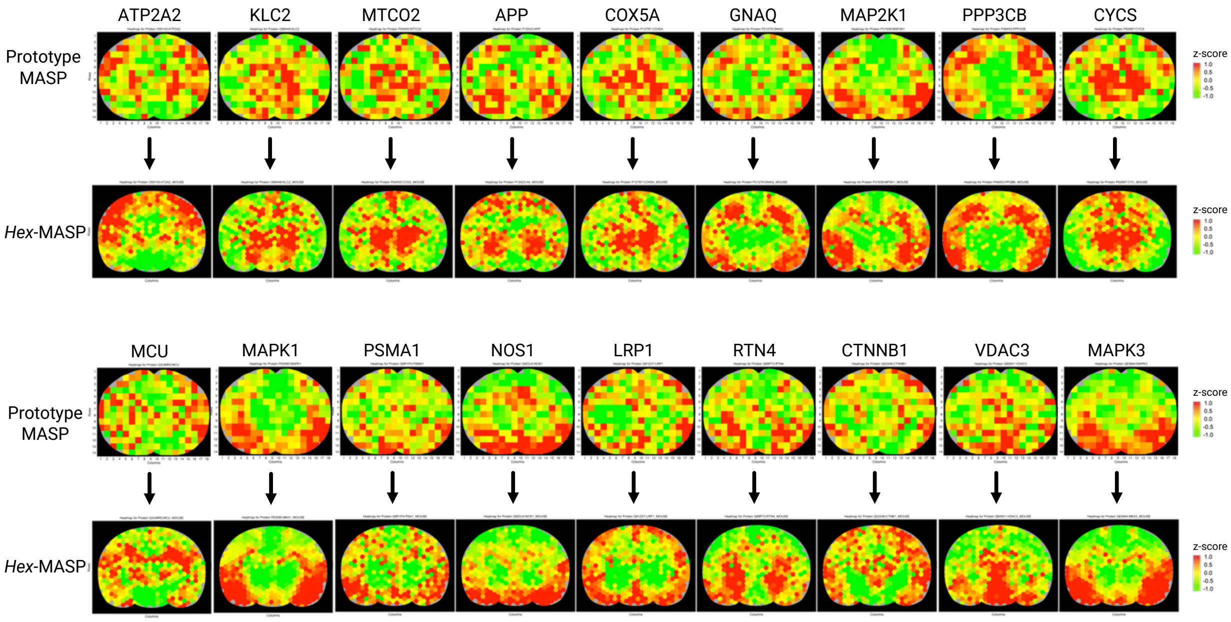

**Supplementary Figure 5. Comparison of the spatial distribution maps of key proteins involved in the KEGG pathway of Alzheimer's Disease acquired by *Hex*-MASP and previous MASP.**

The z-score color scale is from -1.0 (green) to 1.0 (red).

#### Supplementary Results and Discussions

##### Comparison of quantitative accuracy and precision of TMTpro high-throughput mode and IonStar using a benchmark sample set

Isobaric labeling using TMTpro 16-plex enables the simultaneous measurement of peptide abundances in up to 16 samples within a single experiment. Many studies have applied TMT-based approaches for proteome analysis, showing good accuracy, sensitivity, and improved intragroup reproducibility compared with label-free methods[1]. However, to achieve deep proteomic coverage, TMT-based methods often require extensive fractionation (24-48 fractions) of the pooled mixture, especially for tissue samples, which can result in equal or greater instrument time than label-free methods.

Here, our purpose is to take full advantage of the high throughput of TMTpro and combine it with our trapping-nano-LC-MS system to increase throughput without substantially compromising of the proteome depth. Because our system can accommodate complex samples at high protein loading, we anticipate that fewer fractions (*e.g.,* only four) could yield comparable proteome coverage to label-free methods, while enabling at least a fourfold increase in analytical throughput.

To comprehensively assess this approach, we first optimized the LC-MS methods for TMTpro-16plex. We used a set of three-proteome benchmark samples [2] to evaluate the accuracy, precision, and data normalization method of TMTpro-based quantification, in comparison with the IonStar method. Our objectives were to evaluate: i) The proteomic depth achievable in unfractionated vs. fractionated TMTpro samples, ii) The accuracy and precision of TMTpro-based quantification, and iii) The effectiveness of intra- and inter-batch normalization. To prepare these benchmark samples, small, variable amounts of *E. coli* and yeast digests (mimicking altered proteins; “true positives”) were spiked into a large, constant background of mouse digest (representing unaltered proteins; “true negatives”). Five groups were generated with *E. coli* protein ratios relative to Group A at 1×, 1.5×, 2×, 2.5×, and 3× (Groups A-E), as detailed in **Supplementary Table 4**. Each group included three replicates. TMTpro channels were assigned as follows:

- 126: Reference pooled sample
- 127N: Sample A, Replicate 1
- 127C: Sample B, Replicate 1
- 128N: Sample C, Replicate 1
- 128C: Sample D, Replicate 1
- 129N: Sample E, Replicate 1
- 129C: Sample A, Replicate 2
- 130N: Sample B, Replicate 2
- 130C: Sample C, Replicate 2
- 131N: Sample D, Replicate 2
- 131C: Sample E, Replicate 2
- 132N: Sample A, Replicate 3
- 132C: Sample B, Replicate 3
- 133N: Sample C, Replicate 3
- 133C: Sample D, Replicate 3
- 134: Sample E, Replicate 3

##### Identification of proper inter-batch normalization method for TMTpro to improve quantification performance

During these evaluations, we observed a minor ratio compression for *E. coli* proteins, as well as higher inter-batch CV% for TMTpro-labeled samples compared to the label-free workflow. While the ratio compression is acceptable, the batch effect must be addressed. Therefore, here we identified a proper normalization method to address the batch effects.

The strategy for TMT inter-batch normalization was evaluated and performed based on the previous publications[3,4]. We set aside one TMTpro channel in each labeling batch as a common reference. This reference channel contained an aliquot of a pooled sample (composed of 10-20% of randomly selected biological samples). Here we leveraged two well-accepted normalization methods. The first strategy subtracts the log2 transformed intensity measured in the reference channel from the log2 transformed intensity of the same protein measured in other channels within the same batch, to normalize the batch effects. The final step centers the average protein intensities of each batch to zero, following a previously published approach[3]. The second strategy, known as internal reference scaling[4], calculates a scaling factor for each labeling batch and adjusting the protein intensities by the batch-specific scaling factor. Here we incorporated the above methods into the pipeline. Briefly: i) Compute the log2 ratio of each sample-channel intensity to its reference-channel intensity within the same batch, then center the median log2 ratio at zero. ii) Reverse log-transform these ratios to obtain linear protein ratios. iii) Multiply by the average intensity of the reference channels across all batches, producing fully normalized protein intensities. We assessed this combined approach by comparing the intra-group CV% of proteins measured across two labeling batches, before and after normalization, and also compared them to replicates measured within a single batch. Results showed that inter-batch variation was nearly eliminated. The median intra-group CV% dropped from above 20% to below 3% (**Supplementary Figure 2c**), a level comparable to that of samples measured within the same batch (**Figure 2b**). Thus, our combined strategy substantially reduced TMTpro batch-to-batch variation and was subsequently applied to the ICV-dosed mouse brain spatial proteomics study.

#### Supplementary Experimental Procedure

##### Materials and Reagents

Acetonitrile, acetone, ammonium bicarbonate, formic acid, methanol, ethanol, sodium dodecyl sulfate (SDS), and sodium chloride were purchased from Fisher Scientific (MA, USA). Micro-BCA protein-assay kit, the TMTpro 16-plex labeling kit, and the Pierce™ high-pH reversed-phase peptide-fractionation kit were obtained from Thermo Fisher Scientific (CA, USA). Sodium deoxycholate (SDC), IGEPAL CA-630, iodoacetamide, proteomics-grade trypsin, cOmplete™ Mini EDTA-free protease-inhibitor cocktail, hydroxylamine, and Evans-Blue dye came from Sigma-Aldrich (MO, USA). Dithiothreitol (DTT) was sourced from Cytiva (MA, USA), and Tris-base from MP Biomedicals (OH, USA). Other materials: Protein LoBind tubes (Eppendorf, Hamburg, Germany), 1 M HEPES buffer (Santa Cruz Biotechnology, TX, USA), and Sera-Mag SpeedBeads (GE Healthcare/Cytiva, IL, USA). SYLGARD™ 184 silicone elastomer (Dow, MI, USA) and HTM140v2 resin (EnvisionTEC, Germany).

AB095 is the courtesy of AbbVie and an anti-PCSK9 antibody obtained via research prescription from University at Buffalo, NY, USA.

#### Automatic SP3 on KingFisher Flex in µSE-SP3

Preparation of KingFisher plates was performed using the volumes and solvents indicated below, with modifications based on a previous publication.[5]

First, the Tip Comb was set on the Tip Plate. Then, in 96-well deep-well plates, the following reagents were dispensed into each processing lane: i) 50 µL of SP3 bead mix (Beads Plate); ii) 100 µL of protein sample (Sample Plate); iii) 200 µL of 80% ethanol (three Wash Plates). After a pause, 50 µL of freshly prepared digestion buffer (trypsin in 50 mM HEPES, pH 8.0) was added to the Elution Plates.

After reagent loading, the Tip Plate, Beads Plate, Sample Plate, and the three Wash Plates were placed onto the KingFisher Flex system, and the SP3 script was launched. When the script paused for digestion-buffer addition, the trypsin buffer was immediately added to the Elution Plate. Subsequently, the Elution Plate was placed back onto the KingFisher Flex system to continue digestion.

#### TMTpro 16 plex labeling and off-line high-pH reversed-phase fractionation

All location-specific micro-specimens were randomized into twenty-nine TMT batches, each containing peptides from fifteen micro-specimens plus one pooled reference. The reference aliquot was always tagged with channel 126; the remaining samples were distributed across channels 127N through 134N. Labeling employed a 1:1.5 (protein: reagent, w/w) ratio and proceeded for one hour at room temperature in darkness. The reaction was quenched with 1 µL of 5 % hydroxylamine and left for fifteen minutes, also in darkness. All sixteen channels of a batch were pooled and dried in a SpeedVac, re-suspended in 0.1 % trifluoroacetic acid (300 µL), and fractionated off-line with the Pierce high-pH reversed-phase kit. Eight fractions were collected and concatenated (fractions 1+5, 2+6, 3+7, 4+8) to yield four final fractions, which were dried again with SpeedVac and resuspended in 0.1 % formic acid containing 2 % acetonitrile for LC-MS analysis.

#### Protein identification and quantification

Raw data were submitted to the Proteome Discoverer software (v3.0, Thermo Scientific, USA), using the Spectrum Files node. The SEQUEST node was set up to search data against the UniProt (Swiss-Prot) reviewed mouse database (a total of 17129 entries, augmented with AB095 and Anti-PCSK9 sequences). Parameters specified trypsin specificity with up to two missed cleavages, carbamidomethylation (static) on cysteine, TMTpro tags on lysine and peptide N-termini (static), and oxidation of methionine plus protein-N acetylation (dynamic). Reporter ion integration was carried out with 20 ppm tolerance and the most confident centroid was set as the integration method. Percolator was employed with the identification threshold of 1% false discovery rate (FDR) on both protein and peptide levels. After integration of the reporter-ion intensities into protein intensities, the latter was log2-transformed. For normalization, within each batch, the log2 value of the reference channel was subtracted from those of all other channels. Following this, the normalized log2 intensities were back-transformed to linear space and multiplied by the grand-mean reference intensity, yielding the abundance values for each protein in each spatially resolved micro-specimen. More details about the normalization method are in the **Supplementary Results and Discussions.**
